# Cancer models and statistical analysis of age distribution of cancers

**DOI:** 10.1101/2021.03.22.436464

**Authors:** Alexandr N. Tetearing

**Affiliations:** Saint-Petersburg State University

## Abstract

In this paper, mathematical mutational models of the age distribution of cancers are obtained. These are two models – a simple model and a complex model, which takes into account the growth of the cell population and the transmission of mutations to daughter cells. Using the resulting formulas, we approximated real age-specific cancer incidence datasets in women (colon, lung, mammary, stomach) and men (colon, lung, prostate, stomach). We estimated parameters such as the average number of mutations (per cell per unit of time) and number of mutations required for cancer to occur.

The number of mutations averaged (over four types of cancer) required for cancer to occur is 5.5 (mutations per cell for women) and 6.25 (mutations per cell for men) for the complex mutational model.

As an alternative to mutational models, we also consider the model of delayed carcinogenic event.

## 1. Introduction

One of the hallmarks of a cancer cell is a large number of hereditary mutations. In the work of Lawrence Loeb [1], the number of mutations is estimated at a huge number of 10,000 - 100,000 per cell. But this number only applies to long-lived progressive tumours, in which most of the mutations could arise after the degeneration of a normal cell into a cancerous one.

But how many mutations are the minimum required for the formation of a cancer cell?

Alfred Knudson was the first to try to answer this question in 1971. Knudson investigated retinoblastoma, an ocular tumour that occurs in childhood [2]. Based on the available data on 48 cases of retinoblastoma and using the Poisson distribution (mathematical probability distribution of a random event), Knudson concluded that the average number of mutations leading to cancer is three mutations per cell. And the first mutation is a hereditary mutation passed on to a child from a parent.

the question of the minimum required number of cellular mutations leading to malignant transformation remains open.

In this work, mathematical formulas are obtained that describe the age distribution of cancer in a group of patients or in a cell population. When changing the parameter *z* in the formula (this is the number of mutations required for a cancerous transformation of a cell), the curve describing the age distribution of cancer in the population, changes its shape. Comparing theoretical curves with real (obtained in practice) age distributions, we can choose the value of *z* parameter so that the theoretical distribution most closely matches the practical age distribution of cancer.

Thus, we can approximately find out the average minimum number of mutations required for a cancerous cell transformation (if the used mathematical model accurately describes the process of mutation formation).

## Part I Cancer models

### 2. Simple mutational model

#### 2.1. Mathematical model

This simple mathematical model is based on the assumption that the age distribution of cancer is independent of the growth function of the anatomical organ (tissue). This model is an approximate model, since it does not take into account the shape of the growth function of the cell population.

In a simple mutation model, we consider a group of *N*_*P*_ people. Each person from the group has a certain anatomical organ (tissue), which consists of separate identical cells. At the initial moment of time *t* = 0, the cell population of the organ consists of one cell. Over time, the number of cells increases in accordance with the organ growth function *N*(*t*).

All cells are constantly exposed to destructive external influence. As a result of external influence, cells receive mutations. The average number of mutations in a cell per unit time is constant and equal to *a*. The average number of mutations in a cell in a short time *dt* is the product of *a* and *dt*. The average number of mutations in a cell population of *N* cells in a short time *dt* is *N a dt*.

We assume that, with a certain number of mutations accumulated by the cell (we denote this number as *z*), the cell (in any case) is transformed into a cancer cell (perhaps this does not happen immediately, but only after a long time).

Our primary task is to calculate the number of cells in a population with different numbers of mutations, and the number of registered cancer patients in this group.

The damaging effect on the cell population begins at the time *t* = 0. At the time *t* = 0 all cells do not have mutations, and the number of cells with mutations is zero.

Take a look at Fig. 2.1. The figure schematically shows the development of cell populations in a group of 27 people. In each individual person, the growth of an organ (cell population) occurs in accordance with the growth function *N*(*t*). This function is shown as a bar chart on the left side of Figure 2.1.

**Figure 2.1.**
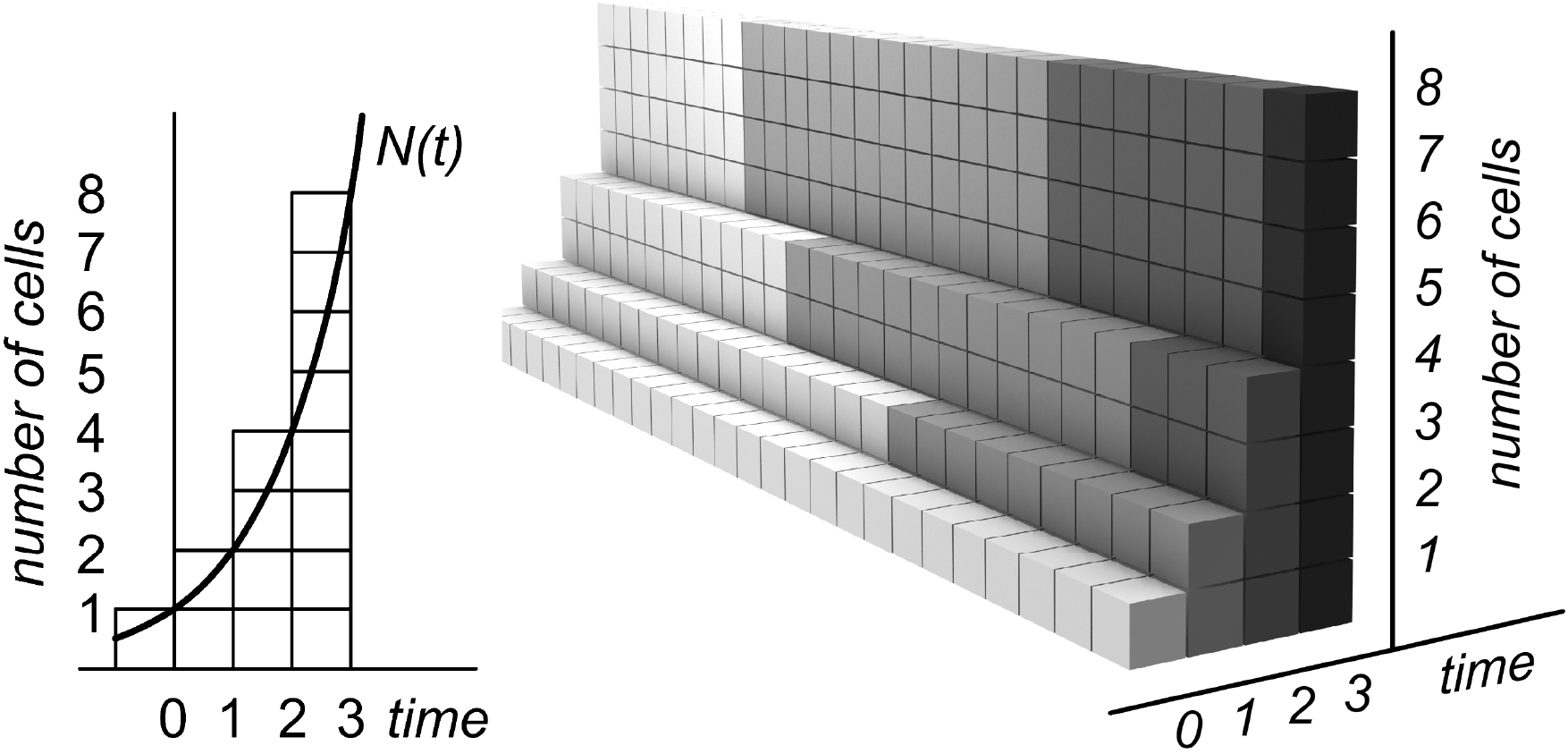
Scheme for obtaining mutations in a group of 27 people with the same functions *N*(*t*) of population growth.

We consider the state of the cell population discreetly – at regular intervals *dt*. In this case, *dt* = 1.

At the moment of time *t* = 0, the organ (cellular tissue) consists of one cell. An organ consists of two cells at time *t* = 1, four cells at time *t* = 2, and eight cells at time *t* = 3. This growth function has been chosen solely for example. In reality, the function may have other parameters.

Thus, at the moment of time *t* = 0, in a group of 27 people there are 27 separate cells (27 people have 27 identical organs, and each organ contains one cell). These are 27 light-colored cells in the first row.

By the time *t* = 1, the number of cells has doubled, and the first mutation has occurred in some of the cells. We consider (for example) that a third of all cells undergo mutation per unit time. That is, in one cell per unit of time, on average, 0.33 mutations occur (in a group of three cells, one cell receives a mutation). 18 cells that have received a mutation at the time interval [0; 1] are shown (in two rows) in light gray. In total, at the moment of time *t* = 1, the figure shows 54 cells (36 of them have no mutations).

For this model, we believe that mutations are inherited from the parent cell to the daughter cells.

By the time *t* = 2, the number of cells will double again – up to 108 (4 rows of 27 cells), and some of the cells will undergo mutation. One third of cells with one mutation will receive the second mutation (cells with two mutations are shown in dark gray). One third of cells that do not have mutations will receive the first mutation. Thus, 12 cells (four rows of 3 cells) by the time *t* = 2 will have two mutations (dark gray cells); 48 cells (4 rows of 12 cells) will each have one mutation (light gray cells).

By the time *t* = 3, the number of cells will double again – up to 216 cells (8 rows of 27 cells), and some of the cells will mutate again. A third of cells with two mutations will receive a third mutation (cells with three mutations are shown in black). A third of cells with one mutation will receive the second mutation (cells with two mutations are shown in dark gray). One third of cells that do not have mutations will receive the first mutation. Thus, by the time *t* = 3, 8 cells (8 rows of 1 cell) will have three mutations (black cells in the figure); 48 cells (8 rows of 6 cells) will have two mutations (dark gray cells), 96 cells (8 rows of 12 cells) will each have one mutation (light gray cells); 64 cells (8 rows of 8 cells) will not have mutations (these are the lightest cells in the picture).

Each vertical column in Fig. 2.1 consists of cells of the same color (these are cells with the same number of mutations). Therefore, we can determine the number of people in whose anatomical organ there are cells with a certain number of mutations by looking only at the cells of the lower layer. This means that in this mathematical model, the function of the number of people in a group who have (for example) one mutation does not depend on the growth function of the cell population (the growth function of cell tissue).

In fact, the described scheme for obtaining mutations does not correspond to reality and is a great simplification. For example, in our model, at time *t* = 1, nine people have cells with one mutation – each of nine people has two cells with one mutation. In reality, mutations in a group are distributed more evenly, and it is more likely that 18 people will receive one mutated cell each.

A simplified scheme for obtaining mutations allows us to disregard the shape of the growth function of the cell population *N*(*t*). A more complex and more realistic model will be discussed in the following sections.

Looking at the bottom layer of cells in Fig. 2.1, we can draw the functions *N*1_*P*_ (*t*), *N*2_*P*_ (*t*), and *N*3_*P*_(*t*). The *N*1_*P*_ (*t*) function describes the number of people who have cells with one or more mutations. The *N*2_*P*_ (*t*) function describes the number of people who have cells with two or more mutations.

The *N*3_*P*_ (*t*) function describes the number of people who have cells with three or more mutations, etc. The *P* index in the function name means that the function describes the number of people, not the number of cells.

The functions *N*1_*P*_ (*t*), *N*2_*P*_ (*t*) and *N*3_*P*_ (*t*) are shown in Fig. 2.2 in the form of bar charts. These functions correspond to Figure 2.1.

**Figure 2.2.**
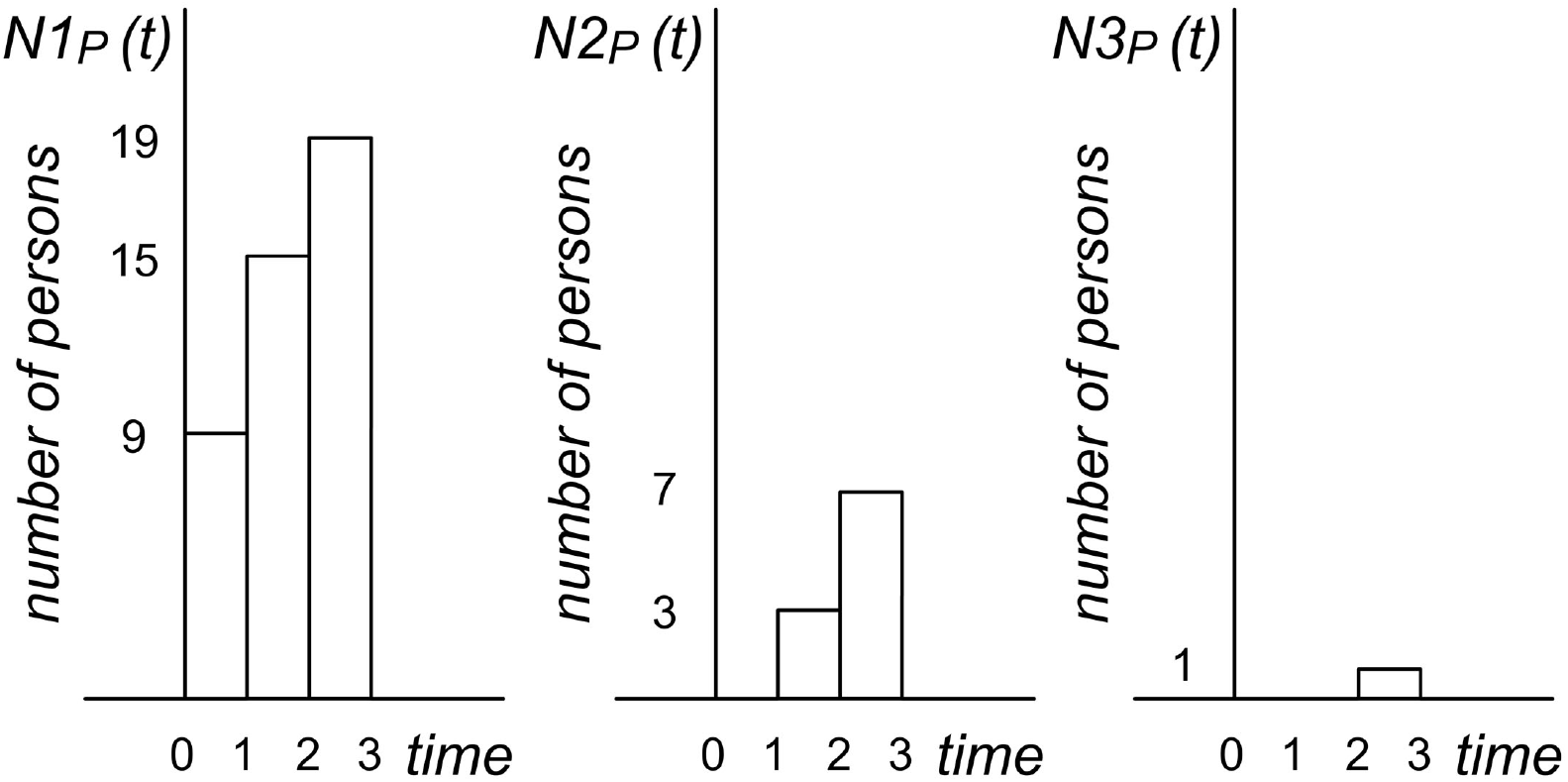
Function *N*1_*P*_ (*t*) – the number of people who have cells with one or more mutations. Function *N*2_*P*_ (*t*) – the number of people who have cells with two or more mutations. Function *N*3_*P*_ (*t*) – the number of people who have cells with three or more mutations. The functions correspond to Fig. 2.1.

The functions shown in Fig. 2.2 are not smooth functions. They are depicted as steps. To get smoother (and more accurate) functions, the time interval *dt* should tend to zero (we record the state of the cell population at regular intervals *dt*). In Fig. 2.1 and 2.2 this interval is equal to one.

Consider the above functions for an arbitrary growth function of the cell population. The number of people in the group under consideration is *N*_*P*_.

At time *t* = *dt*, the number of people with mutated cells is:

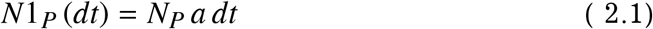

Here *a* is the average number of mutations that a cell receives per unit of time.

The number of people without mutations *N*0_*P*_ is:

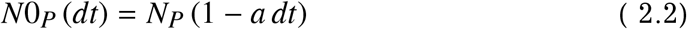

At time *t* = 2*dt* the number of people without mutations *N*0_*P*_ is:

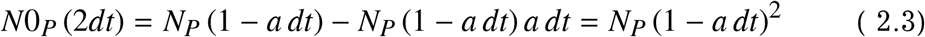

At time *t* = 3*dt* the number of people without mutations *N*0_*P*_ is:

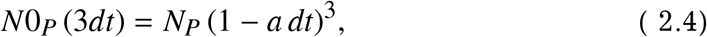

etc.

Thus, the number of people without mutations at time *t* = *T* is the product of *N*_*P*_ (group size) and a definite multiplical of the negative constant *a* on the interval [0; *T*]:

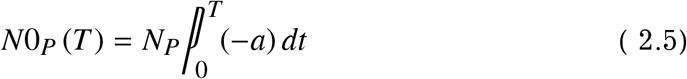

The definite multiplical of the function *f* (*t*) is the limit of the infinite product:

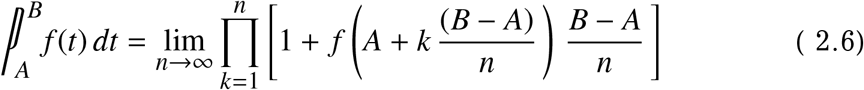

In accordance with the basic formula of multiplical calculus, the definite multiplical of the function *f* (*t*) is the difference of the antiderivatives *F*(*t*) at the ends of the interval:

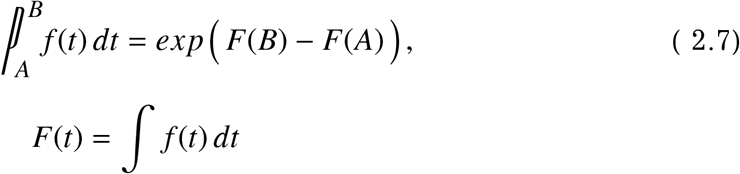

The derivation of the formula is given in [3]. In our case, since *a* is a constant:

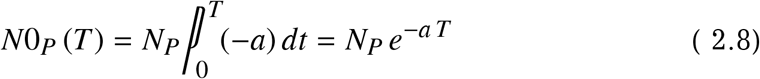

The number of people who have at the time *t* cells with one mutation (or with a large number of mutations) is:

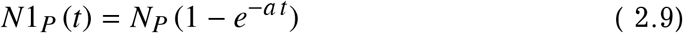

The graph of the function *N*1_*P*_ (*t*) is shown in Fig. 2.3. We used the following parameters to plot the graph: *N*_*P*_ = 1000; *a* = 0.05.

**Figure 2.3.**
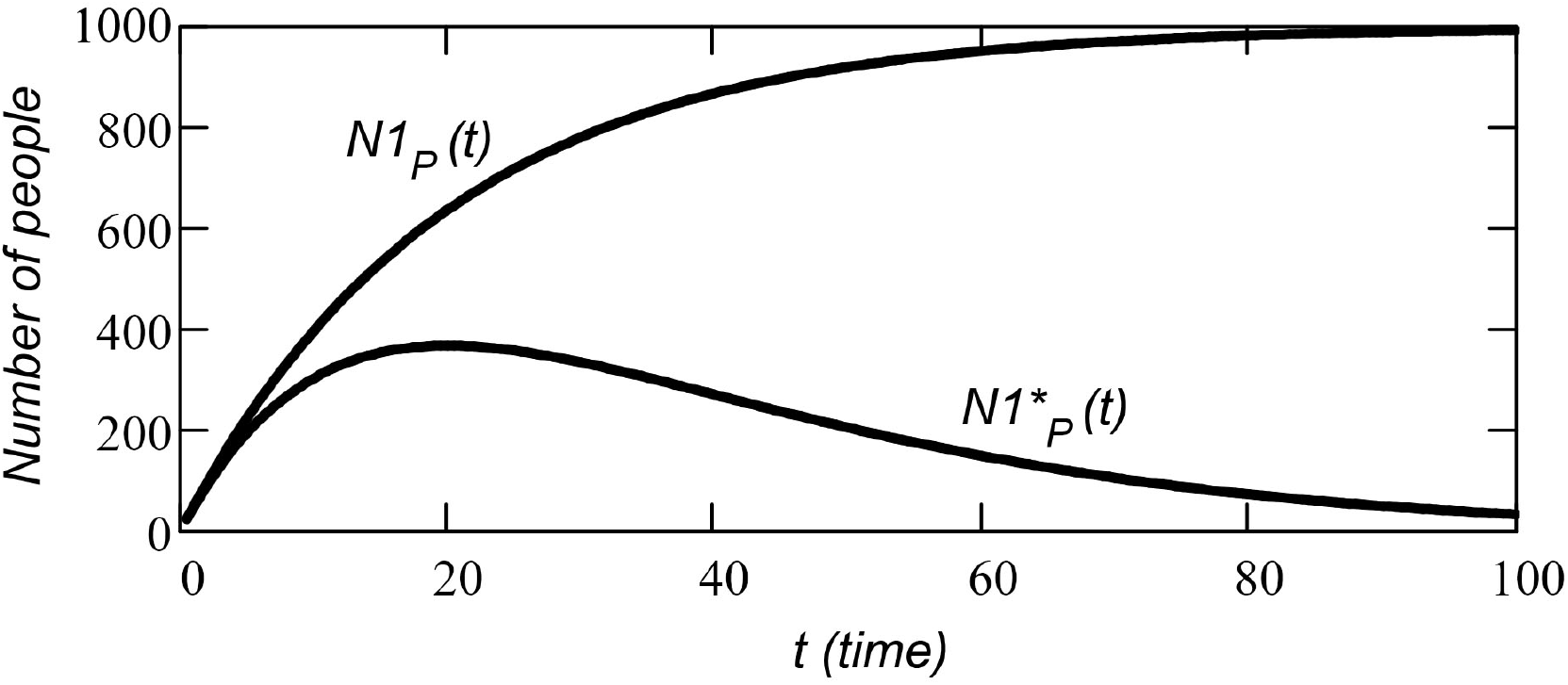
The number of people who have (at the time *t*) the cells with one mutation (or more) – function *N*1_*P*_ (*t*) (upper curve in the figure). The number of people who have cells with only one mutation – function 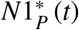 (bottom curve).

Now let’s find the number 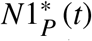 of people in the group who at time *t* = *T* have cells with only one mutation (no more and no less) and who do not have cells with the number of mutations exceeding one.

People who have cells with only one mutation are formed from people who did not have mutations, the number of which is described by the function *N*0_*P*_(*t*).

At each time interval *dt* (starting from the moment *t* = *dt*), there are two processes that change the number of people in the group who have cells with one mutation.On the one hand, the available number of people with one mutation decreases due to the fact that some people receive the second mutation per cell. On the other hand, the number of people who have cells with one mutation increases due to the fact that some people without mutations receive one mutation. (on each interval *dt* this is the increment of the function *N*1_*P*_ (*t*) on the interval *dt*).

Since the function *N*1_*P*_ (*t*) is a function that changes over time, we cannot simply use a formula in the form (2.5) to find 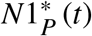. Instead, we need to calculate the product of *dN*1_*P*_ (the increment of the function *N*1_*P*_ (*t*)) and a definite multiplical on each *dt* interval. Let’s look at Fig. 2.4.

**Figure 2.4.**
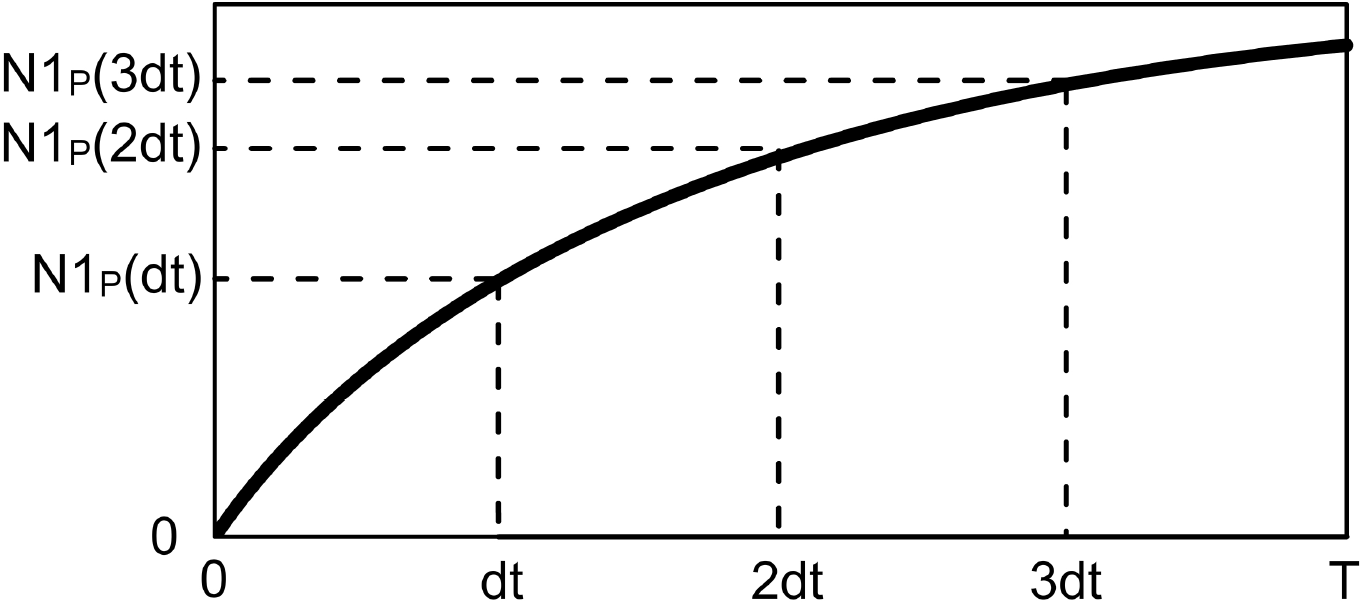
To the calculation of the function *N*2_*P*_ (*t*) (the number of people in a group who have cells with two mutations).

The known function *N*1_*P*_ (*t*) on the interval [0; *T*] is bounded, positive and monotonically increasing. Divide the interval [0; *T*] into small segments *dt*. The number of these segments is *n* = *T* /*dt*. In this case, *n* = 4.

Then the function 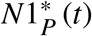 is the sum:

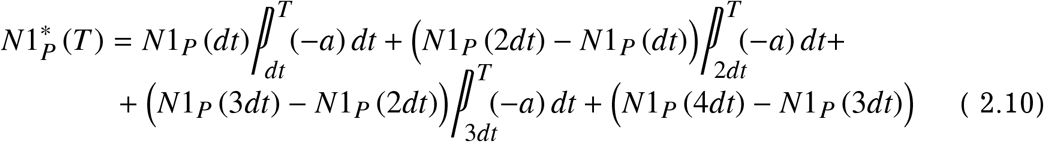

or:

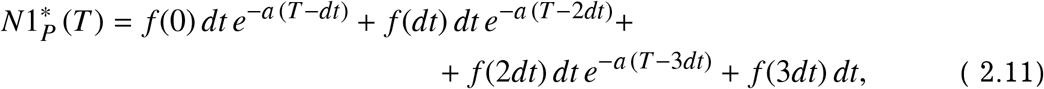

where *f* (*t*) is the time derivative of the function *N*1_*P*_ (*t*):

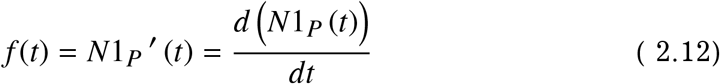

If *dt* is infinitesimal (for *n longrightarrow in f ty*), then it is true:

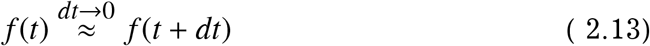

That is to say, 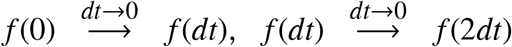 etc. Moreover, 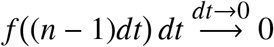. Then:

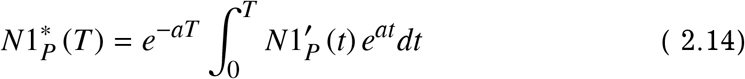

We will use this formula in the future, but, in this case:

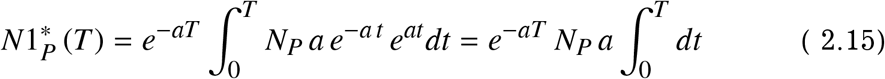

Thus, the number of people, who have cells with only one mutation (who do not have cells with a number of mutations exceeding one), changes over time as a function of 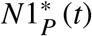:

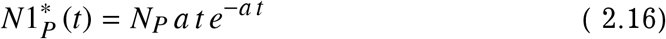

We can calculate the function 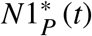 (the number of people who have cells with at most one mutation) in a different way. To do this, we need to solve a differential equation:

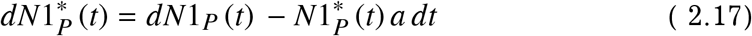

The left side of the equation is the change in the number of people who have the cells with one mutation (on the time interval *dt*). The first term on the right side is the number of people that received the cells with one mutation in *dt* time (these people are formed from people that have not previously been mutated).

The second (negative) term on the right is the number of people that received the cells with a second mutation in the *dt* interval (these people are formed from people who previously had cells with only one mutation).

Dividing the equation by *dt* we have:

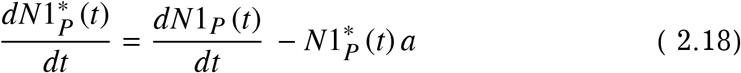

We introduce the notation:

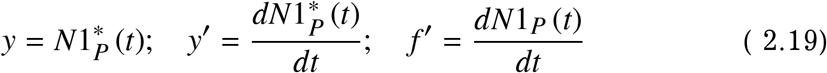

Linear non-homogeneous differential equation:

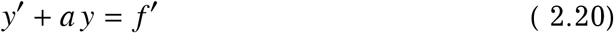

is solved by the substitution *y* = *uv*.

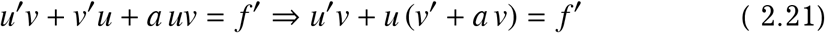

When the expression in parentheses is zero: (*v*′ + *a v*) = 0, we get the following expression for *v*:

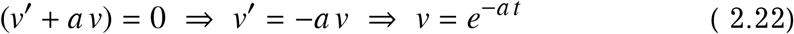

The resulting expression is substituted into equation (2.21):

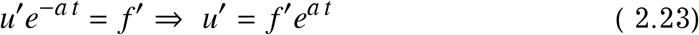

Integrating both sides of the equation, we get:

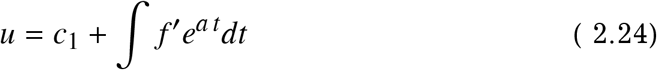

Then we substitute the functions *u* and *v* into the expression *y* = *uv* and return to the notation (2.19):

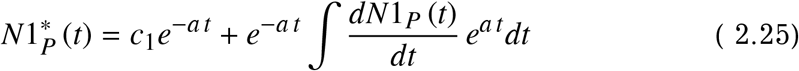

Substitution the function *N*1(*t*) into this equation gives (see Eq. 2.9):

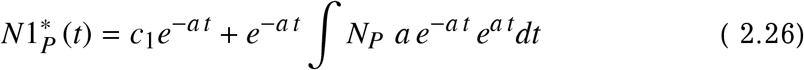

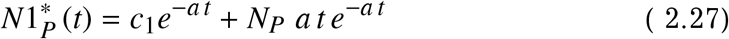

To find the integration constant *c*_1_, we substitute *t* = 0 into this equation:

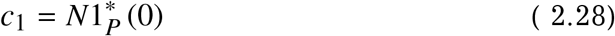

In our case, 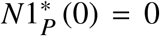. As a result, we get:

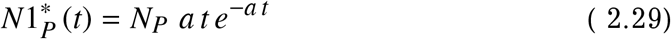

So, we obtained formula (2.16) in a different way.

The number of people *N*2_*P*_ with two mutations per cell (or more) is the difference of the functions *N*1_*P*_ (*t*) and 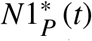:

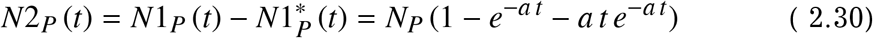

The graph of the function *N*2_*P*_ (*t*) is shown in Fig. 2.5 (upper curve).

**Figure 2.5.**
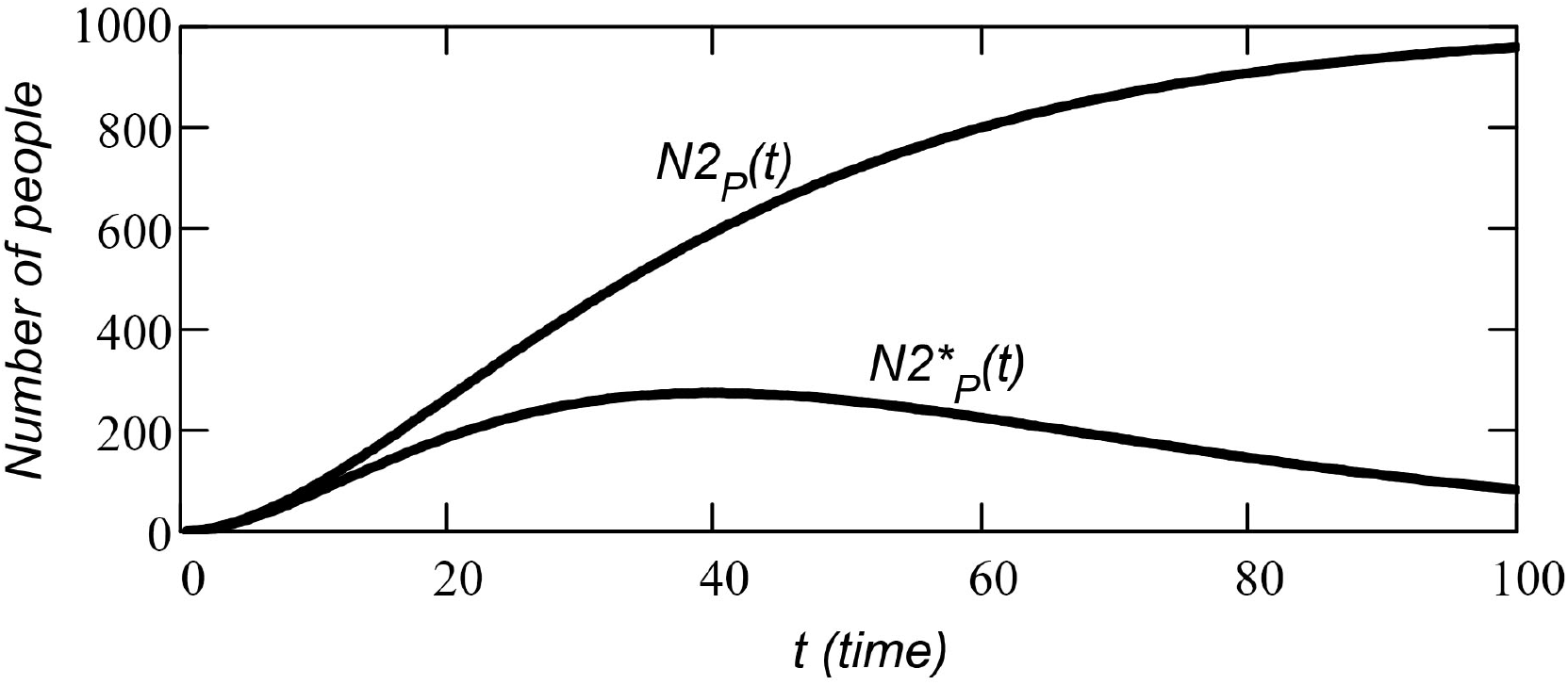
The number of people who have cells with two (or more) mutations – the function *N*2_*P*_ (*t*) (upper curve in the figure). The number of people who have cells with only two mutations (who do not have cells with fewer or more mutations) – the function 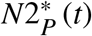 (bottom curve).

To find the number of people 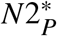 who have cells with only two mutations (no less, no more), we will use the formula (2.14) obtained earlier. Now we need to substitute the function *N*2_*P*_ (*t*) into this formula instead of the function *N*1_*P*_ (*t*).

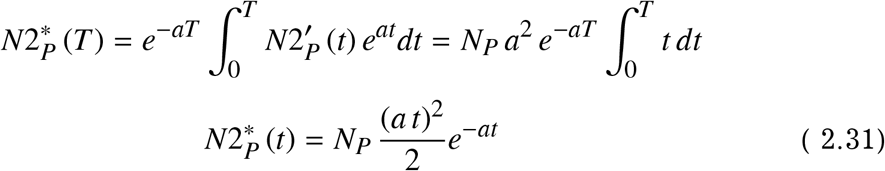

The graph of the function 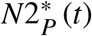 is shown in Fig. 2.5 (bottom curve).

If we subtract the function 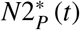 (number of people who have cells with exactly two mutations) from the function *N*2_*P*_ (*t*) (number of people who have cells with two or more mutations), we get the function *N*3_*P*_ (*t*) (the number of people who have cells with three or more mutations).

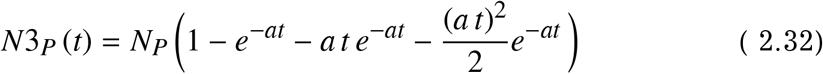

To find the function 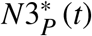 (number of people who have cells with exactly three mutations), we again use formula (2.14). In this time, we need to substitute the function *N*3_*P*_ (*t*) into the formula instead of the function *N*1_*P*_ (*t*):

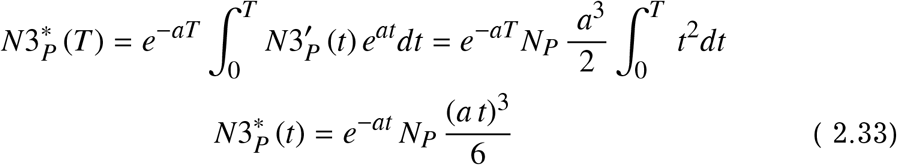

The graphs of the functions *N*3_*P*_ (*t*) and 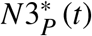 are shown in Fig. 2.6.

**Figure 2.6.**
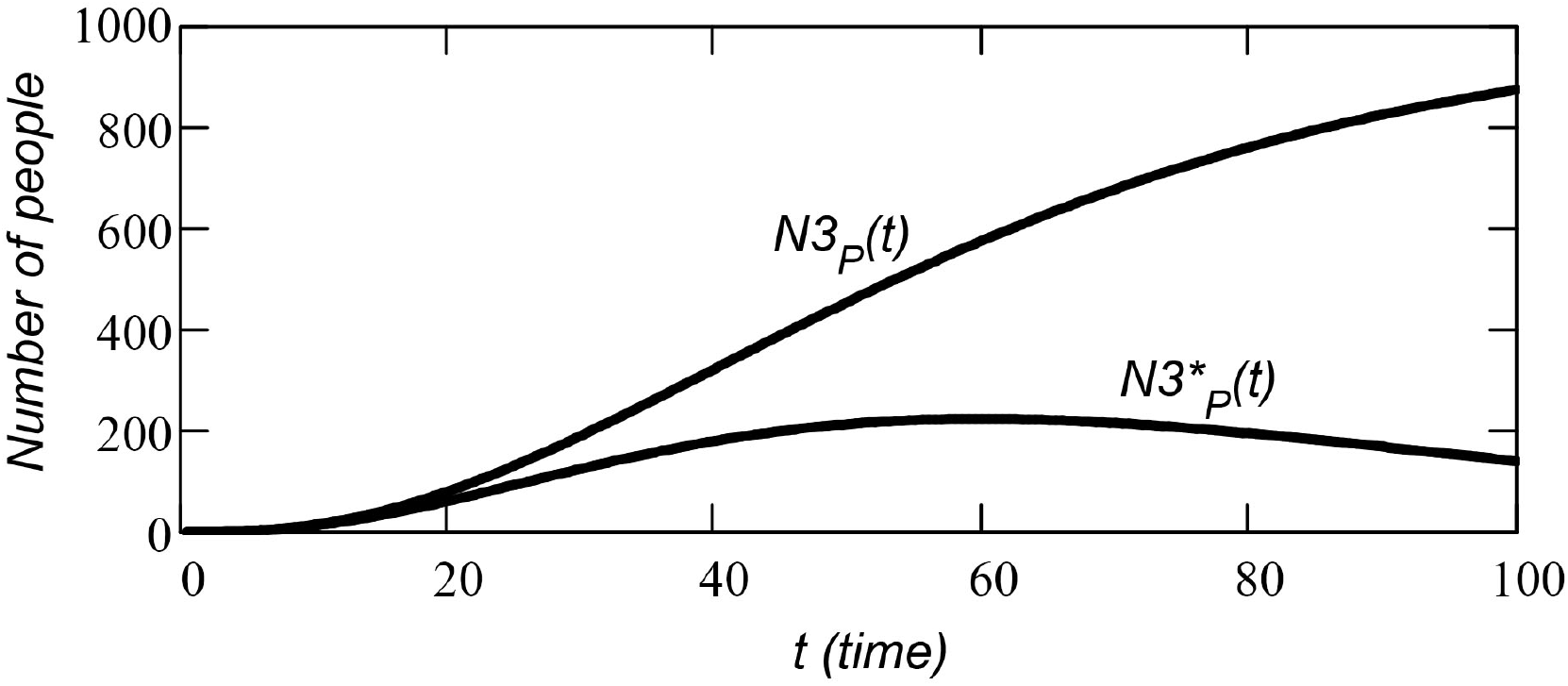
The number of people who have cells with three (or more) mutations – the function *N*3_*P*_ (*t*) (upper curve in the figure). The number of people who have cells with exactly three mutations – the function 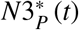 (bottom curve).

Similarly, we find the functions 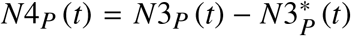 (the number of people who have cells with four mutations or more) and 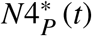 (the number of people who have cells with exactly four mutations).

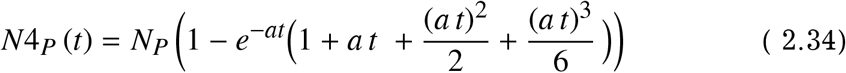

To find the function 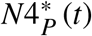 (the number of people who have cells with exactly four mutations) we use formula (2.14). We substitute the function *N*4_*P*_ (*t*) into the formula instead of the function *N*1_*P*_ (*t*):

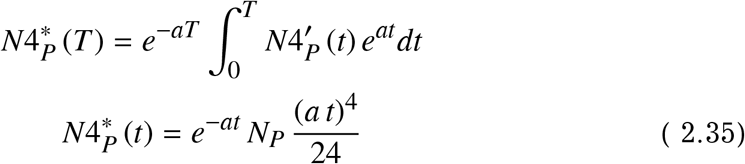

The graphs of the functions *N*4_*P*_ (*t*) and 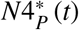 are shown in Fig. 2.7.

**Figure 2.7.**
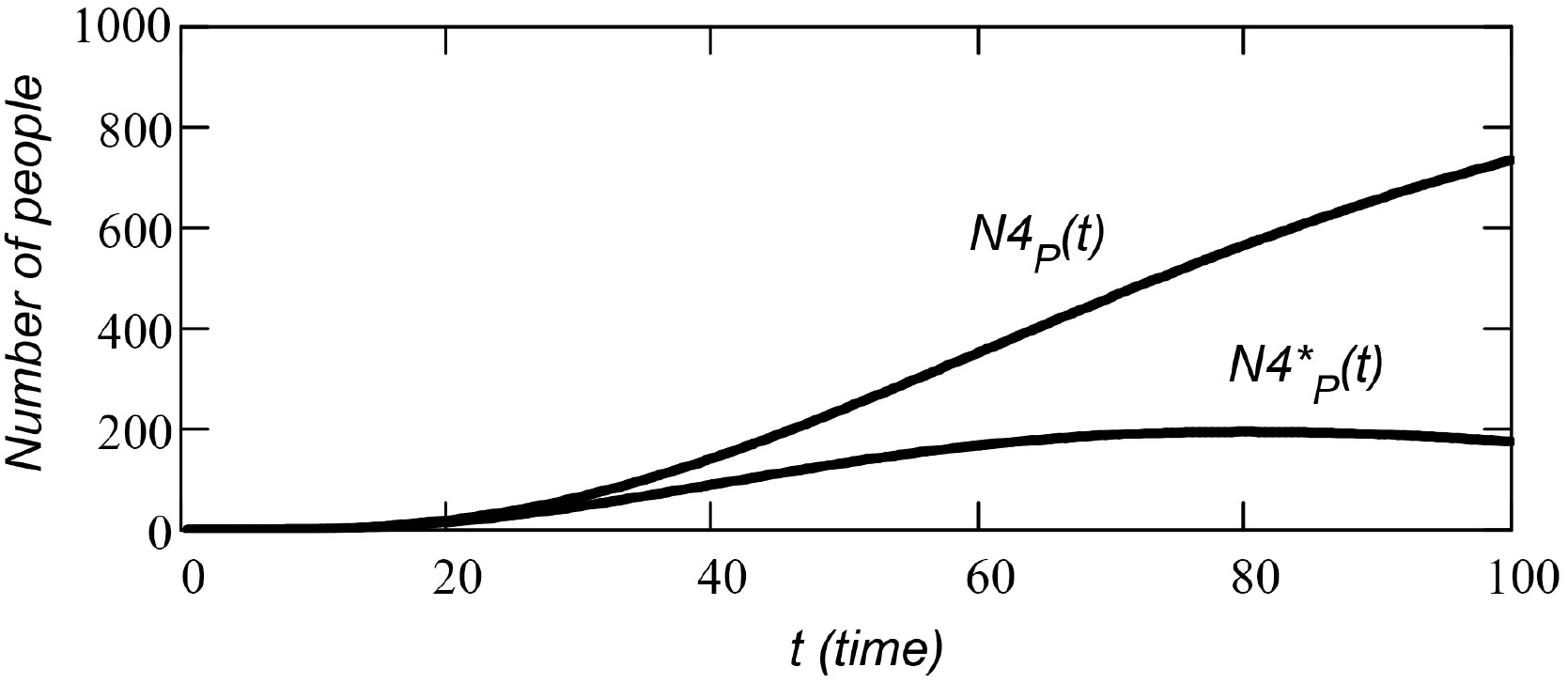
The number of people who have cells with four (or more) mutations – the function *N*4_*P*_ (*t*) (upper curve in the figure). The number of people who have cells with exactly four mutations – the function 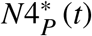 (bottom curve).

Looking at the formulas for functions *N*1_*P*_ (*t*), *N*2_*P*_ (*t*), *N*3_*P*_ (*t*), and *N*4_*P*_ (*t*), it is easy to deduce the general formula for the function *Nz*_*P*_ (*t*), where *z* is the minimum number of mutations in a cell of an anatomical organ in one person.

For example, the designation *N*8_*P*_ (*t*) corresponds to a function that describes the number of people in a group that have cells with the number of mutations from 8 or more.

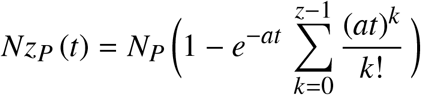

Here *N*_*P*_ is the size of the group.

#### 2.2. Age distribution of cancers

We agreed (this is just for example) that if a cell gets three (or more) mutations, the cell becomes cancerous. To find the age distribution of cancers in the cell population, we need to calculate the derivative of the function *Nz*_*P*_ (*t*) (in our case, this is the function *N*3_*P*_ (*t*)). The function increment on an time interval *dt* is the number of cancers that appeared during this time period.

For example, the age distribution of cancers for the three mutations *D*3(*t*) is:

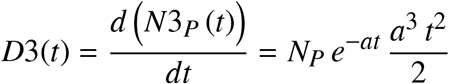

The distribution is shown in Fig. 2.8.

**Figure 2.8.**
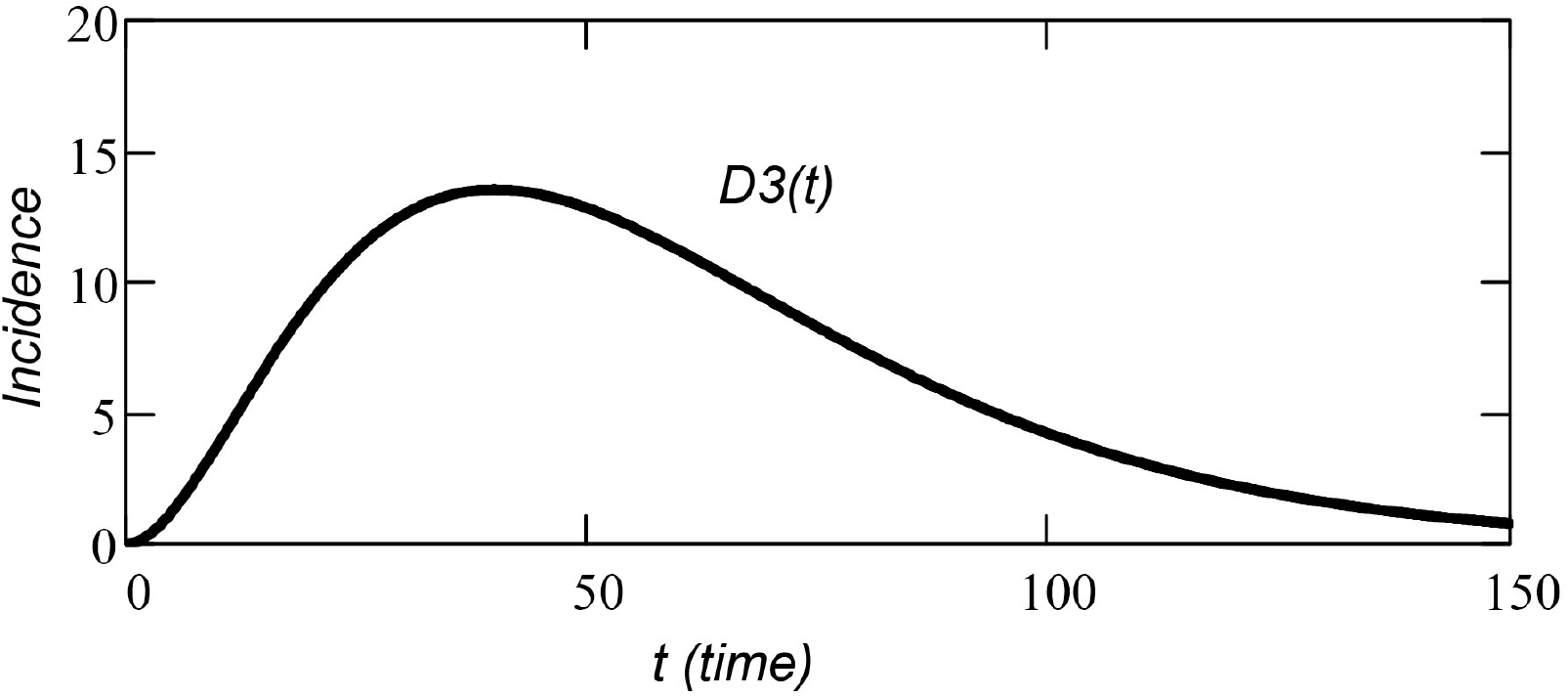
Age distribution of cancers in a group of 1000 people. The case when the disease begins with three mutations in the cell. Simple (rough) mutation model.

The general formula for the distribution of cancers, that arise as a result of a certain number of mutations in a cell, is as follows:

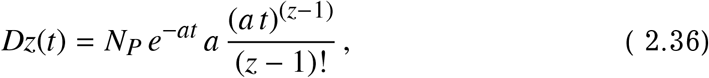

where *z* is the number of mutations in the cell, after which cancer necessarily begins. *N*_*P*_ is the number of people in the considered group.

This is the Erlang distribution known in mathematics [4]. The Danish mathematician Agner Krarup Erlang got this distribution when he worked on the problem of determining the waiting time for a telephone exchange operator.

The graphs of the age distribution for different numbers of mutations are shown in Fig. 2.9. These are the cases when the disease begins with two mutations in the cell (the function *D*2(*t*)), with four mutations in the cell (the function *D*4(*t*)), and with eight mutations in the cell (the function *D*8(*t*)).

**Figure 2.9.**
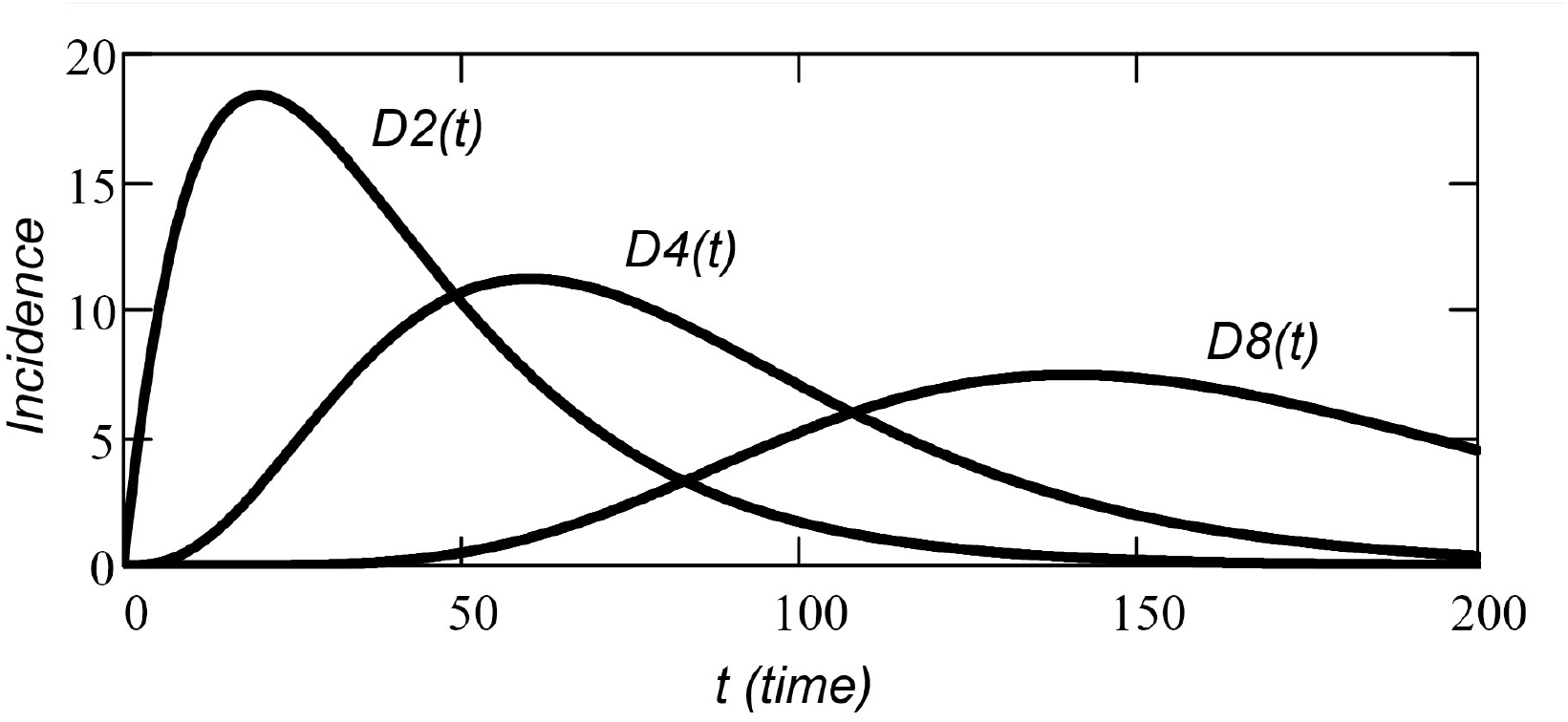
Age distribution of cancers in a group of 1000 people. The cases when the disease begins with two mutations in the cell (the function *D*2(*t*)), with four mutations in the cell (the function *D*4(*t*)), and with eight mutations in the cell (the function *D*8(*t*)).

For all functions considered, the number of people in a group is *N*_0_ = 1000 people, the average number of mutations per cell is *a* = 0.05 mutations per unit of time.

The number of cancers in the group, in the interval [*t*; *t* + *dt*], is the product of the age distribution function *Dz*(*t*) and the length of the interval *dt*.

### 3. Complex mutational model

#### 3.1. Mathematical model

In the previous section, we obtained the age distribution of cancers in a group of people without taking into account the shape of the growth function of the anatomical organ (cell tissue). However, this simplest model, as we have seen, does not correspond to reality. Real cell tissue grows, increasing the number of cells by several orders of magnitude during the growth process, and the changes in growth function can affect the shape of the age distribution.

Consider a cell population consisting of one cell *N*(0) = 1 at the initial moment of time *t* = 0. Here *N*(0) is the number of cells at time *t* = 0.

The growth of the cell population follows the equations:

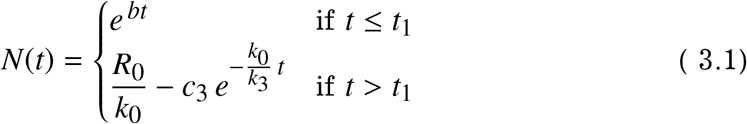

Until the time *t* = *t*_1_, the function is a solution to the equation of unlimited growth – the cell population consumes as much food (energy) resource as it can absorb:

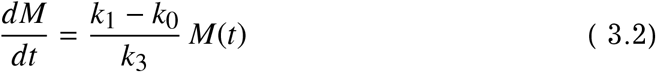

Here *M*(*t*) is the mass of the cell population. To find the size of the population *N*(*t*), we need to divide the mass of the population by the average mass of one cell *m*_0_:

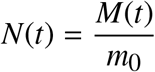

In our case, we assume that the cell mass is equal to one.

After time *t* = *t*_1_, the mass *M*(*t*) of the cell population is a solution to the equation of limited growth – the population consumes a limited (constant) amount of food (energy) resource, because the environment cannot provide more resources:

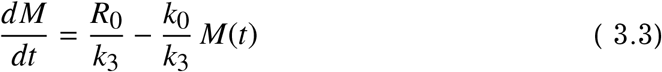

The *R*_0_ constant is the amount of food resource that is available to the population (the amount of resource that the environment provides per unit of time).

According to the equations of population theory [5], only the ratio between the coefficients matters, but not the absolute values of the coefficients *k*_0_, *k*_1_, *k*_3_. That is, the equations do not change if all three coefficients are multiplied or divided by some number. Therefore, it is often believed that the coefficient *k*_1_ is equal to one.

The coefficient *k*_0_ is always less than *k*_1_, otherwise the mass of the population will decrease over time. All the equations and coefficients that characterize the cell population are described in detail in [5].

The constant *b* from the upper equation of the system (3.1) is determined by the expression:

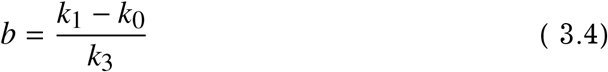

The time *t*_1_ (the time when the cell population goes from the stage of unlimited growth to the stage with limited resource consumption) is determined by the equality:

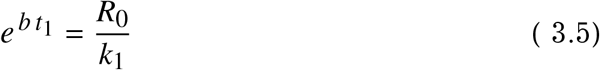

This equality arises from the condition:

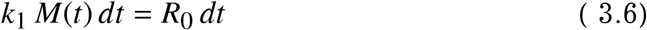

(see Eq. 3.2 and 3.3).

Under this condition, equation (3.2) turns into equation (3.3).

To find the constant *c*_3_ (this is the constant of integration) we solve the equation:

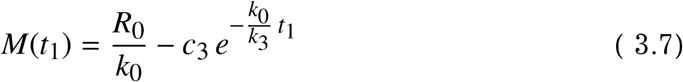

(see System of Equations 3.1).

The development graph of the cell population is shown in Fig. 3.1 (the function *N*(*t*) is the number of cells in the population).

**Figure 3.1.**
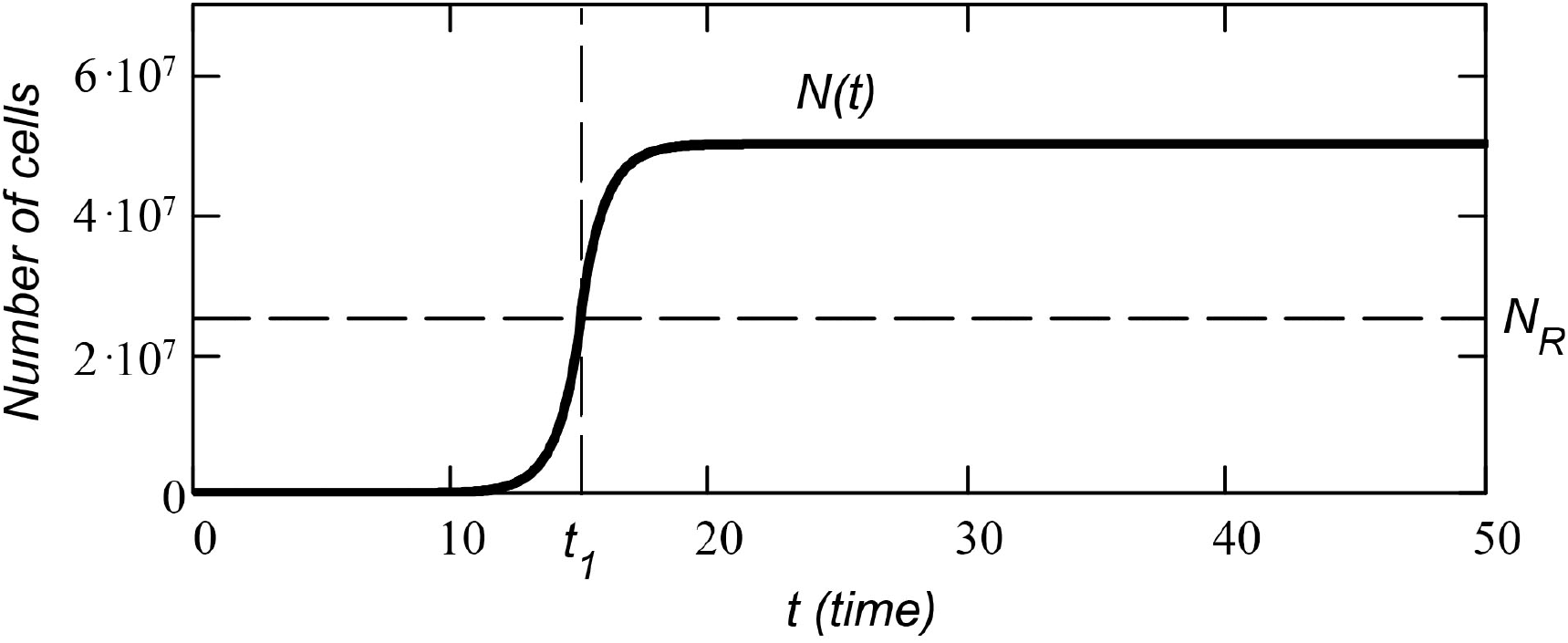
Function *N*(*t*) is the number of cells in the population. The cell population has exponential growth until time *t* = *t*_1_ and limited growth after time *t* = *t*_1_.

The *N*(*t*) function (this is number of cells in the population) demonstrates exponential growth up to the time *t* = *t*_1_ and limited growth after the time *t* = *t*_1_. At the moment *t* = 10, the number of cells in the population is *N*(10) = 8.6 *cdot*10^4^. The population size at the time *t* = *t*_1_ is *N*(*t*_1_) = *N*_*R*_ = 2.5 · 10^7^. With limited growth, the number of cells gradually reaches an asymptotic plateau *N*(∞) = *R*0/*k*_0_ = 5 · 10^7^.

For plotting graph 3.1, we used the following parameters: *R*_0_ = 2.5 · 10^7^; *k*_0_ = 0.5; *k*_1_ = 1; *k*_3_ = 0.44; *t*_1_ = 15.

The number of cells that do not have mutations *N*0 (*t*) is the product of the function *N*0 (*t*) and the definite multiplical of a constant negative number *a* on the interval [0; *T*] (*a* is the average number of mutations per cell per unit of time).

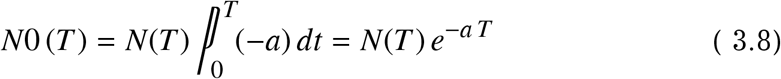

In our case:

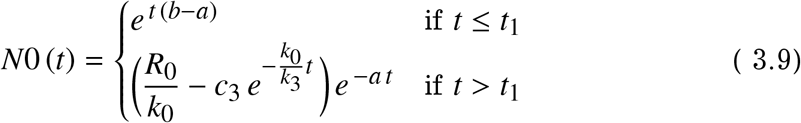

The number of cells with one mutation (or more) *N*1(*t*) is the difference of the functions *N* (*t*) and *N*0 (*t*):

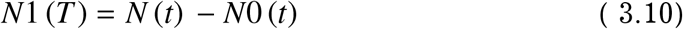

The number of cells with exactly one mutation *N*1^*^(*t*) (no less, no more) is the product of the function *N*1(*t*) and the definite multiplical of a constant negative number *a*:

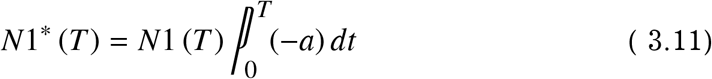

Here *T* and *t* is the same variable. Different notations are used to avoid confusion with the limits of the multiplical.

In our case:

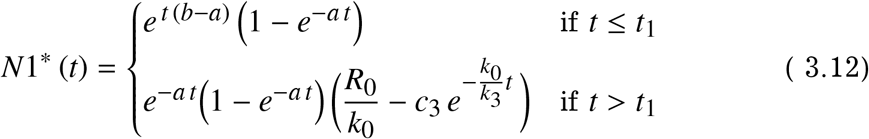

The graphs of the functions *N*1(*t*) (number of cells with one or more mutations) and *N*1^*^(*t*) (number of cells with exactly one mutation) are shown in Fig. 3.2.

**Figure 3.2.**
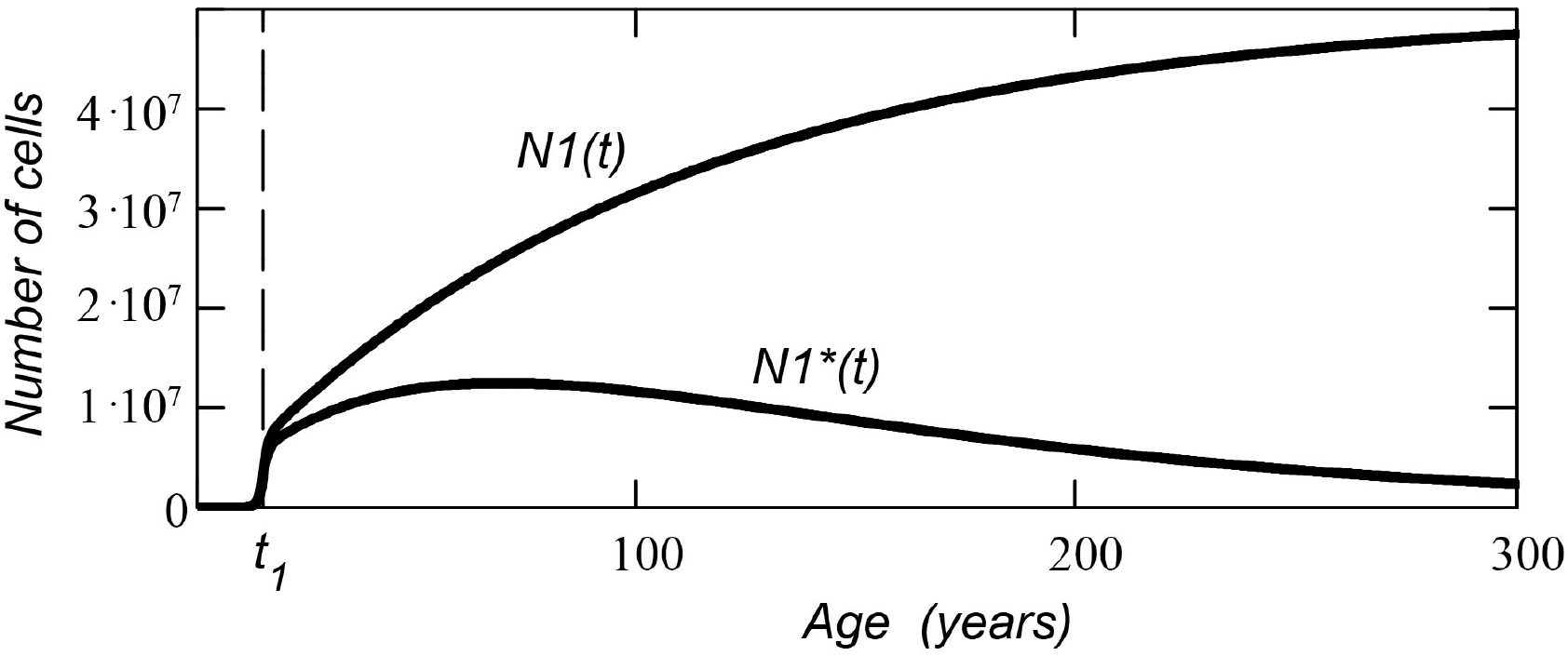
Number of cells with one or more mutations (the function *N*1(*t*) and upper curve in the figure) and number of cells with exactly one mutation (the function *N*1^*^(*t*) and bottom curve).

To find the number of cells with two or more mutations (function *N*2(*t*)), we need to subtract the function *N*1^*^(*t*) from the function *N*1(*t*):

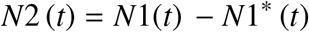

Or:

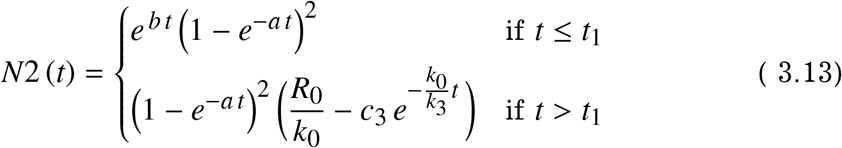

The number of cells with exactly two mutations (the function *N*2^*^(*t*)) is:

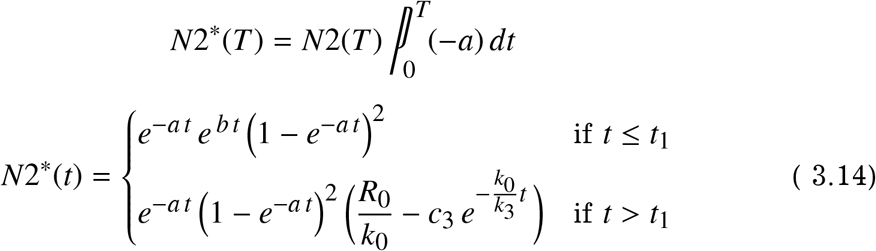

The graphs of the functions *N*2(*t*) (the number of cells with two or more mutations) and *N*2^*^(*t*) (the number of cells with exactly two mutations) are shown in Fig. 3.3.

**Figure 3.3.**
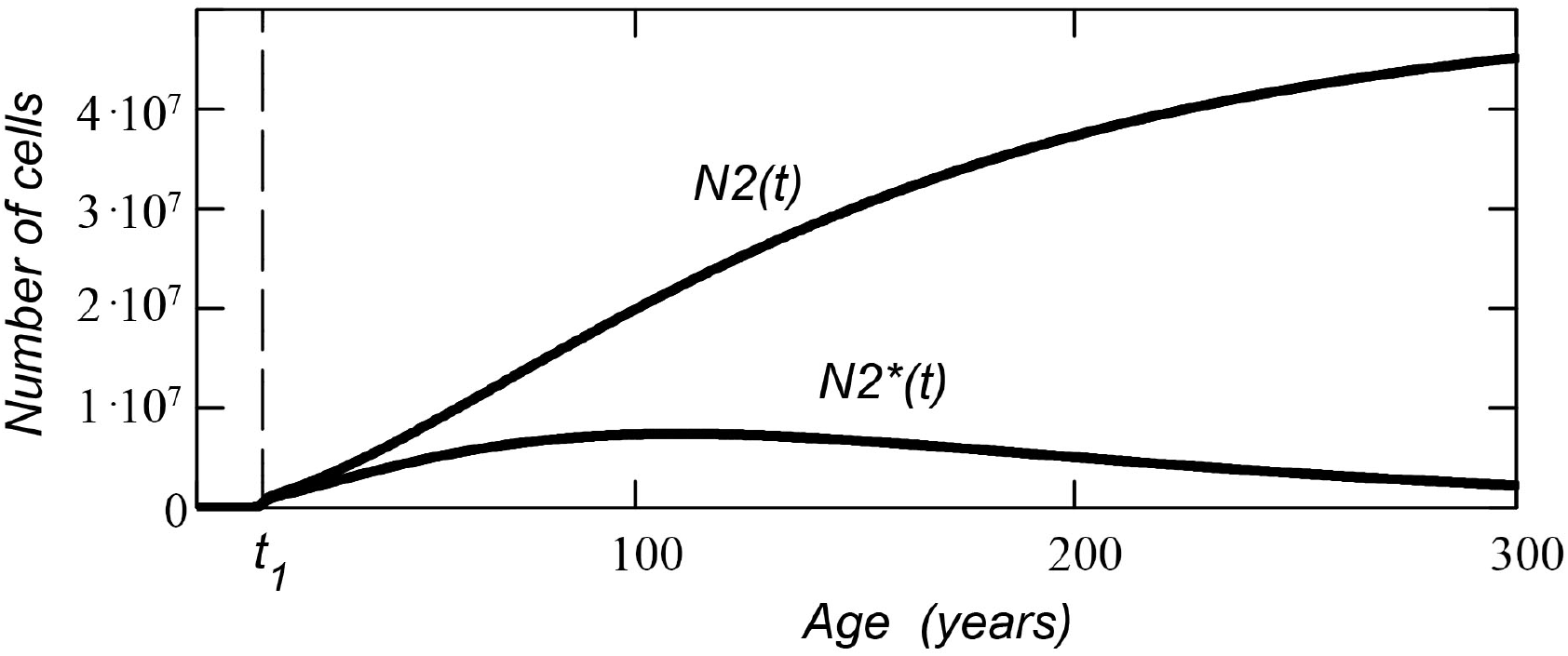
Number of cells with two or more mutations (the function *N*2(*t*) and upper curve in the figure) and number of cells with exactly two mutations (the function *N*2^*^(*t*) and bottom curve).

The formula for the function *N*3(*t*) (the number of cells with three or more mutations) is found in a similar way.

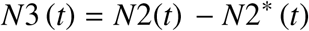

Or:

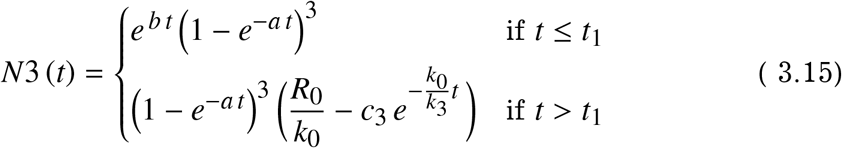

The function *N*3^*^(*t*) (the number of cells with exactly three mutations) is:

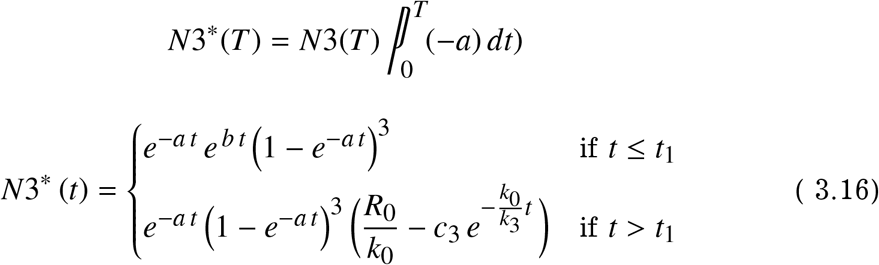

The graphs of the functions *N*3(*t*) (the number of cells with three or more mutations) and *N*3^*^(*t*) (the number of cells with exactly three mutations) are shown in Fig. 3.4.

**Figure 3.4.**
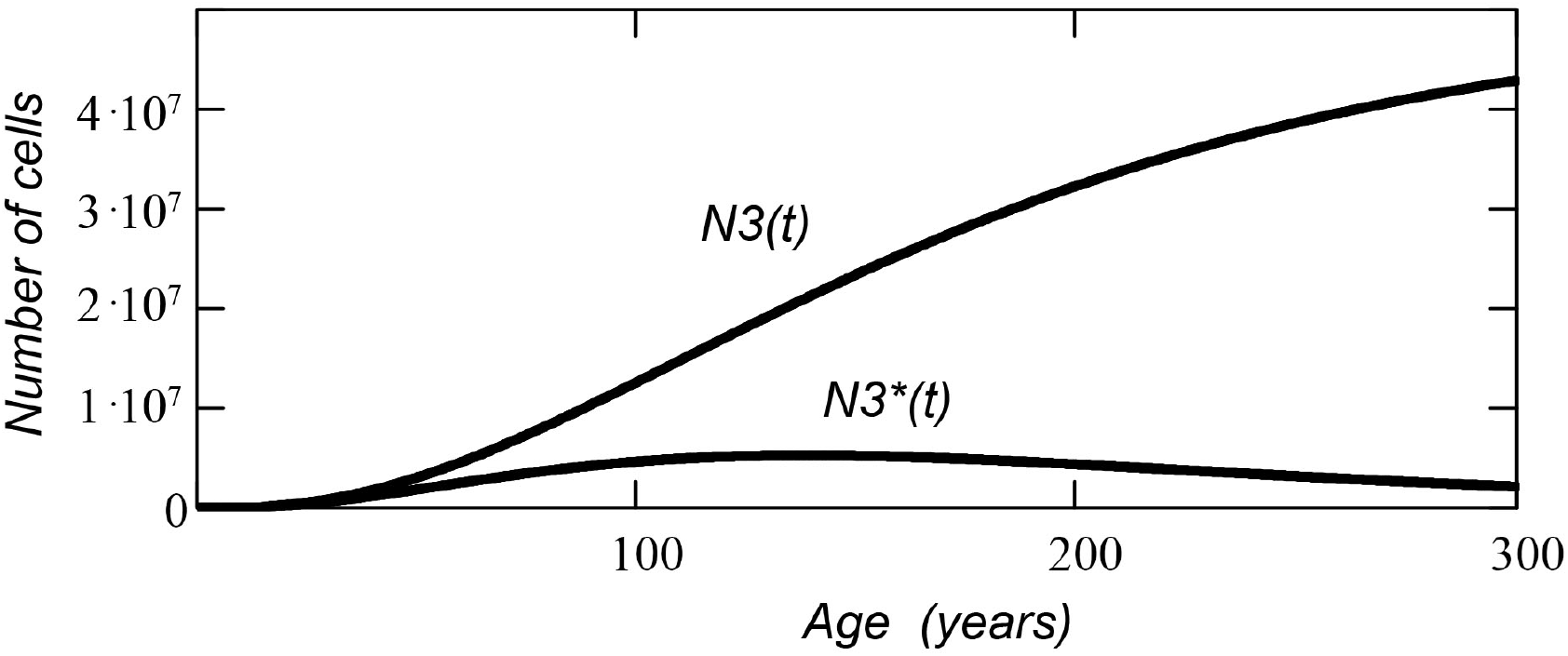
Number of cells with three or more mutations (the function *N*3(*t*) and upper curve in the figure) and number of cells with exactly three mutations (the function *N*3^*^(*t*) and bottom curve).

Looking at these functions, we can derive a general formula for the function *Nz* (*t*) with a certain (minimum specified) number of mutations (where *z* is the number of mutations in the cell):

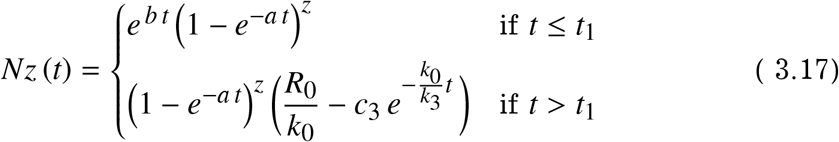

The *Nz* (*t*) function describes the number of cells in which the number of mutations is equal to or greater than *z*. We recall that the time *t* = *t*_1_ is the initial parameter of the cell population, which is determined by formula (3.5).

#### 3.2. Age distribution of cancers

Using the formulas in the previous section, we can get the age distribution of cancers for a group of *N*_*P*_ people.

We assume (for example) that cancer starts after three mutations in one cell. The number of cells with three mutations in a group of *N*_*P*_ people is the function *N*3_*P*_ (*t*):

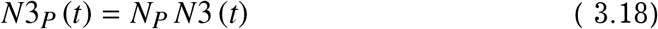

The function *N*3 (*t*) is the number of cells with three mutations (or more) in the cell population of a separate anatomical organ (tissue).

At the initial moment of time, the number of healthy people in the group is *N*_*P*_.

Cells with three mutations are formed from cells with two mutations. In a short time interval *dt* a certain number of cells with three mutations will appear in the population. This number is 5:

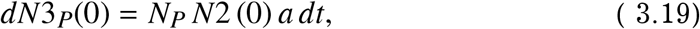

where *N*2 (0) is the number of cells with two mutations in the cell tissue of one person at time *t* = 0. The constant *a* is the average number of mutations that are formed in a cell per unit of time.

If the interval *dt* is small, and the size of the group is large enough, then the number of sick people *dN*3_*P*_ (0) in the group is equal to the number of formed cancer cells.

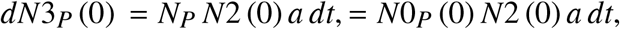

Here *N*0_*P*_ (0) is the number of healthy people in the group at time *t* = 0. On the next interval *dt*, the increase in the number of sick people is:

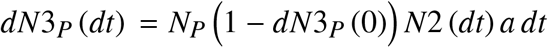

We excluded from the healthy group people who got sick in the previous time interval, so there are no people with three mutations in the remaining group. Cells with three mutations are again formed in an amount proportional to the number of cells with two mutations.

At an arbitrary moment of time *t* the number of healthy people in the group is equal to the product of the initial size of the group *N*_*P*_ and a definite multiplical on the interval [0; *t*] of the product of the function *N*2(*t*) and the parameter *a*, taken with minus sign (see equations 2.6 and 2.7):

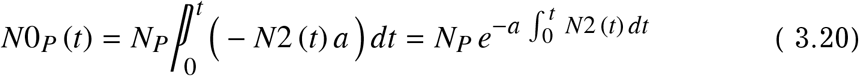

The number of sick people *N*3_*P*_ (*t*) is:

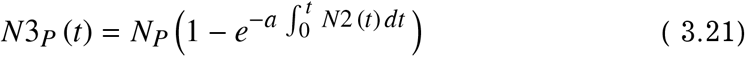

The number of sick people in a group of *N*_*P*_ people is the number of people who have cells with three mutations.

To get the age distribution *D*3(*t*) of cancers in the group, we need to find the derivative of the function *N*3_*P*_ (*t*) :

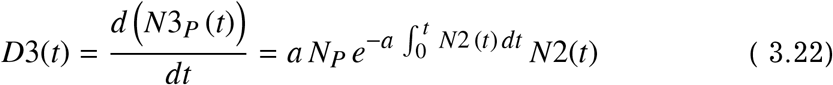

The age distribution *Dz*(*t*) of cancers in a group of *N*_*P*_ – for the case when the number of mutations in a cell required for cancer to occur is z – is as follows:

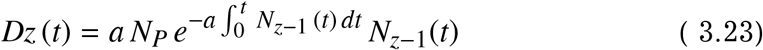

Here the function *N*_*z*−1_(*t*) is the number of cells with the number of mutations (*z* − 1) in the cell tissue of one person. This function is given by the equation:

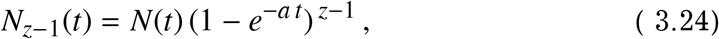

where *N*(*t*) is the number of cells in the cellular tissue of an anatomical organ of one person (this is a function of the growth of cellular tissue – see equations 3.17).

Thus, we can write the formula for the age distribution of cancers *Dz* (*t*) for any arbitrary function *N*(*t*) that describes the growth of the cell population of anatomical tissue.

Substituting another function *N*2(*t*) (the number of cancer-producing cells in the tissue) into the equation, we can find the age distribution for other models of cancer.

The number of cancers that will happen in a group of *N*_*P*_ people in a time interval [*t*; *t* + *dt*], is the product of the value of the age distribution function *Dz* (*t*) by the length of the interval *dt*. Here *z* is the number of cell mutations required for cancer formation.

When using formulas in numerical calculations, we should keep in mind that if the function *N*(*t*) is a piecewise function (as in our case, when the function is given by two equations: one equation before the time *t*_1_, the other equation after the time *t*_1_), then the integral for time *t* > *t*1 must be calculated separately over two segments [0; *t*_1_] and [*t*_1_; *t*]:

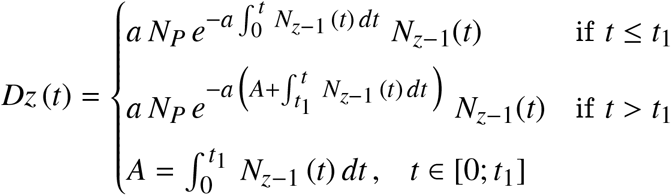

Fig. 3.5 shows the age distributions of cancers for cases where cancer occurs with three, four and five mutations in the cell. The size of the group under consideration is *N*_*P*_ = 1000 people. We used the following parameters for plotting the graphs: *R*_0_ = 5.6 · 10^5^; *k*_0_ = 0.5; *k*_1_ = 1; *k*_3_ = 0.6; *t*_1_ = 15.

**Figure 3.5.**
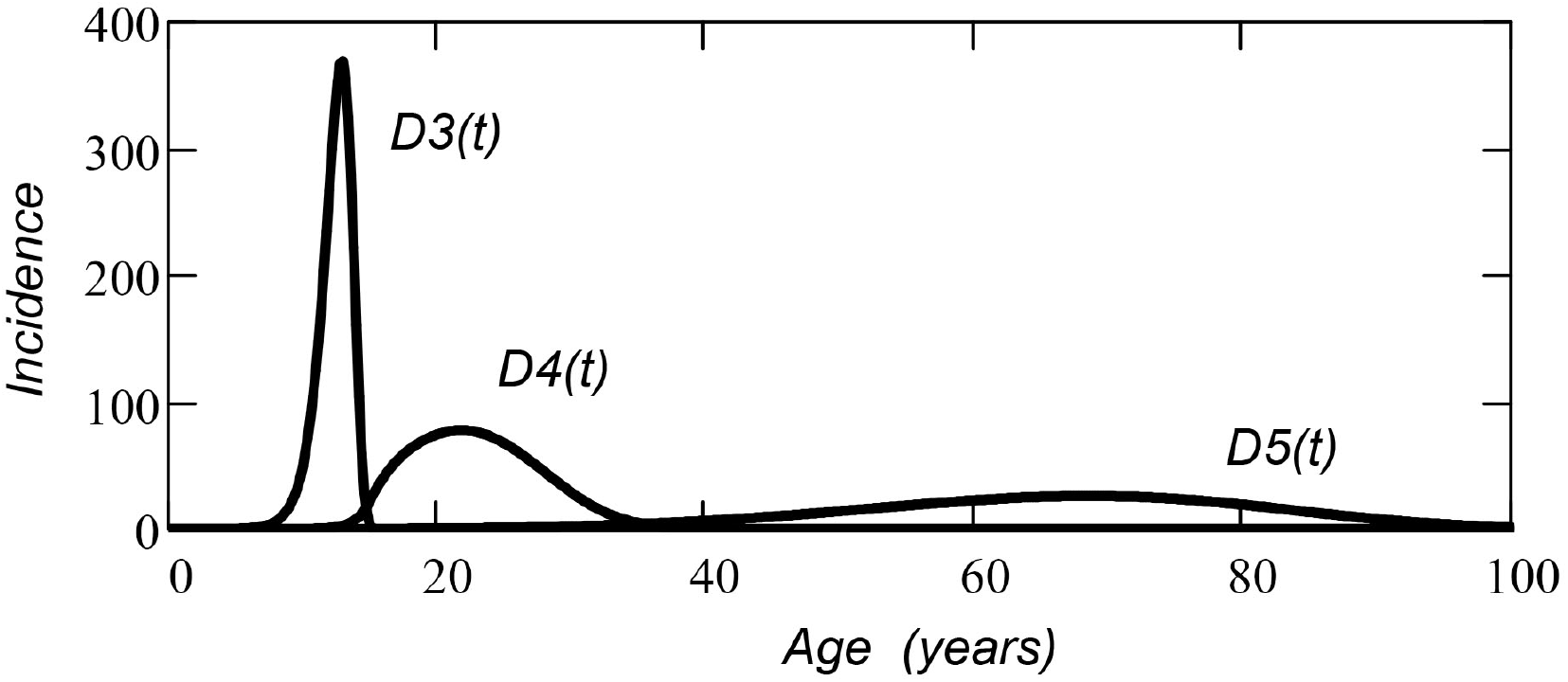
Age distribution of cancers in a group of *N*_*P*_ = 1000 people: with 3 mutations in a cell (the function *D*3(*t*)); with 4 mutations in a cell (the function *D*4(*t*)); with 5 mutations in a cell (the function *D*5(*t*)).

All obtained age distributions must satisfy the normalization condition - the area under the curve *Dz* (*t*) is equal to the size of the group under consideration:

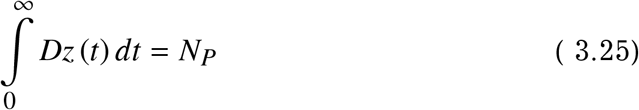

Figure 3.6 shows the age distributions for 4 mutations in a cell for various parameters *a* (average number of mutations in a cell per unit time). Functions are shown for *a* = 100, *a* = 30, *a* = 5.

**Figure 3.6.**
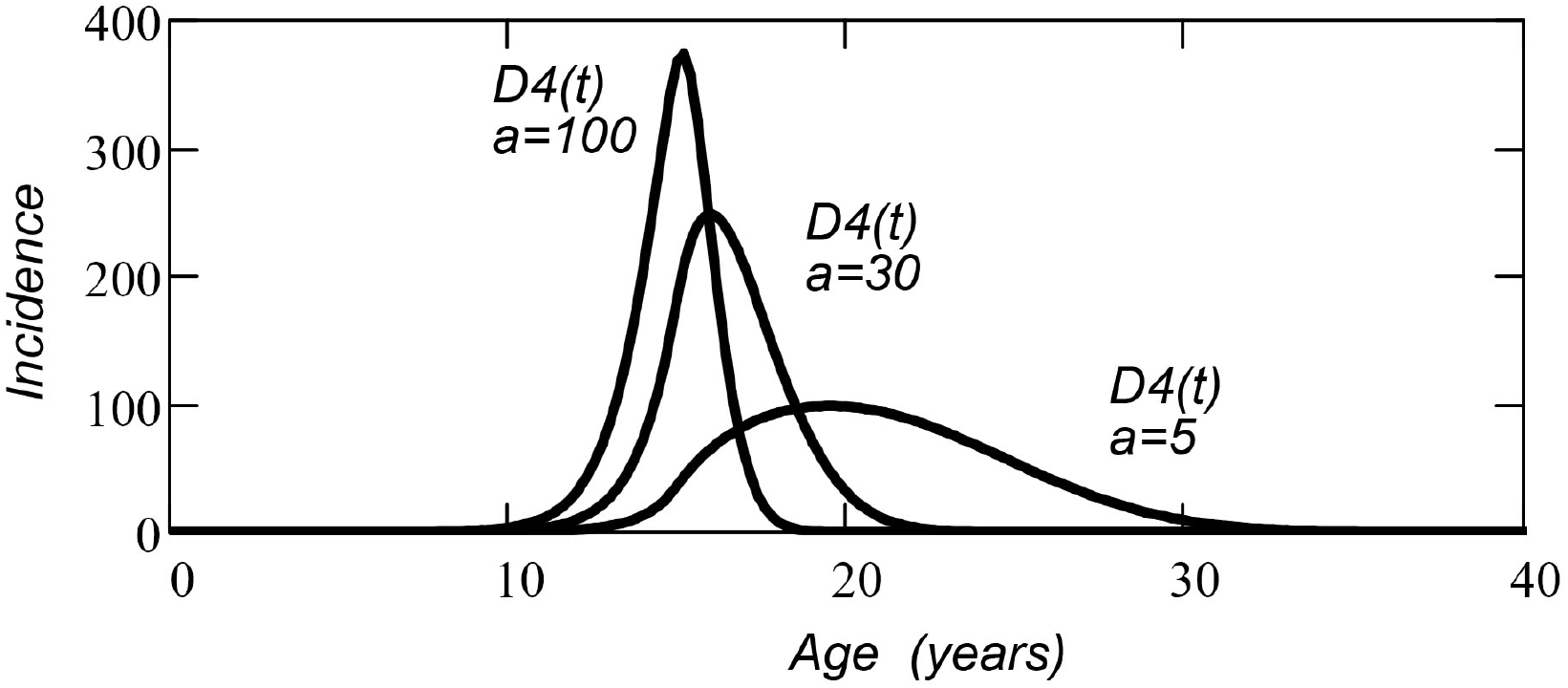
Age distribution of cancers in a group of *N*_*P*_ = 1000 people in the case when cancer forms with 4 mutations in the cell for different parameters *a* (average number of mutations in a cell per unit time).

We can obtain formulas (3.20) and (3.21) in another way. Perhaps for some readers this method will be more understandable.

Let’s say we are considering a group of *N*_*P*_ people. At the initial moment of time *t* = 0, the number of sick people is zero: *N*3_*P*_ (0) = 0. As before (for example only), we believe that cancer in a cell begins when the cell receives three (or more) mutations.

The first cancer patient in the group will appear at the time *t*_1_, when the function *N*3_*P*_ (*t*) is equal to one.

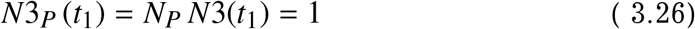

The *N*3 (*t*) function describes the number of cells (in an anatomical organ) that have three (or more) mutations. The function *N*3_*P*_ (*t*) is the number of people who have cells with three (or more) mutations.

After the first cancer patient appears, the number of healthy people in the group will decrease by one. We exclude from consideration people who got the disease (because they have already been taken into account in our statistics), despite the fact that they continue to receive mutations in their cells.

The second cancer patient in a group of (*N*_*P*_ − 1) healthy people appears at the time *t*_2_ when the following condition is met:

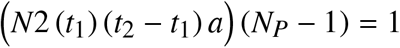

The number of sick people at the time *t*_2_ is 2:

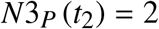

The third cancer patient in a group of (*N*_*P*_ − 2) healthy people appears at the time *t*_3_ when the following condition is met:

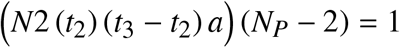

The number of sick people at the time *t*_3_ is 3:

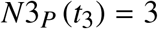

Similarly, we can get a discrete set of values of the function *N*3_*P*_ (*t*_*n*_) = *n* for a discrete set of time points *t*_*n*_. The general equation for *t*_*n*_ is as follows:

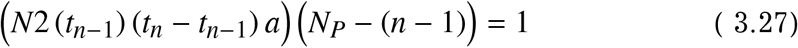

Thus, we get a discrete function *N*3_*P*_ (*t*_*n*_) – that is, a function that is defined only at some points. For practical use, it is more convenient to have a continuous function, which we can get by converting the equation to differential form:

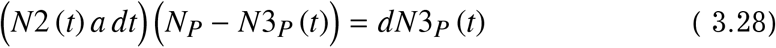

Here *dN*3_*P*_ (*t*) is the increment of the function *N*3_*P*_ (*t*) over a small time interval *dt*.

After separation of variables:

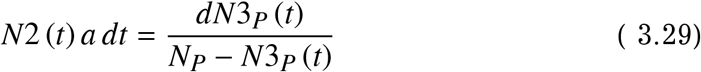

we integrate both sides of the equation:

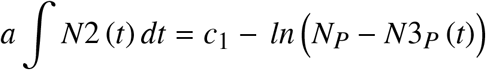

To find the constant *c*_1_, we put the time *t* = 0 into this equation.

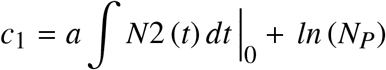

Here ∫*N*2 (*t*) *dt*|_0_ is the value of an antiderivative of the function *N*2 (*t*) at time *t* = 0.

Then:

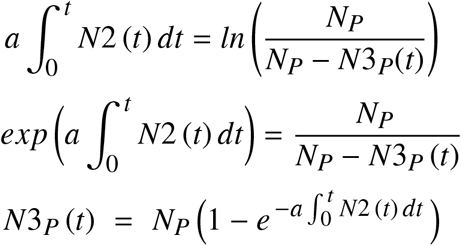

We obtained formula (3.20) in a different way – by composing and solving the differential equation (3.28). The formula describes the increase in the number *N*3_*P*_ (*t*) of cancer patients in a group of *N*_*P*_ people, for the case when the cancer begins with three mutations in the cell.

To obtain the age distribution *D*3(*t*) of cancers in the group, we need (as before) to find the derivative of the function *N*3_*P*_ (*t*) (see Eq. 3.21).

### 4. Model of delayed carcinogenic event

We can use the formulas obtained in the previous section to construct other models of cancer. Let us consider the simplest model of a carcinogenic event with an age shift.

In this model, we assume that the cell becomes cancerous after some single event that has occurred in the cell.

In addition, we assume the presence of an age-related shift – a malignant effect on tissue cells does not begin at age zero, but some time later, several years after birth.

Let the time of the onset of the carcinogenic effect be *T*_*S*_. For the onset of a cancer, one initiating event is enough (what kind of event it is, we do not know yet). The average number of initial events that occur with a biological cell per unit of time is *a*. Until the time *T*_*S*_, there are no cancer cells in the cell tissue. After *T*_*S*_ time, the number of sick people in a group of *N*_*P*_ begins to gradually increase.

We assume that the time *T*_*S*_ is sufficient for the cell tissue to fully form. That is, the growth function of cell tissue *N*(*t*) has reached a horizontal plateau, and the number of cells in the cell tissue does not change over time: *N*(*t*) = *c*_1_.

We can use the previously obtained formula (3.22) to construct the age distribution *D*1(*t*) of cancers in the group. In this case, *z* = 1 (*z* is the number of carcinogenic events in a cell required for cancer to form). Substitute the function *N*(*t*) = *N*_0_ into equation (3.22) instead of the function *N*_*z*−1_:

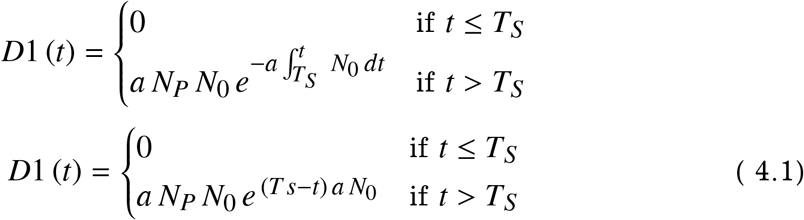

Here *N*_*P*_ is the number of people in the group under consideration; *N*_0_ the number of cells in the cell tissue. In our case (for example), *N*_*P*_ = 1000, mbox *N*_0_ = 2 *cdot*10^6^. The graph of the function *D*1 (*t*) is shown in Fig. 4.1-a. To plot the graph, we used the following parameters: *a* = 8 *cdot*10 ^−8^; *T*_*S*_ = 30.

**Figure 4.1.**
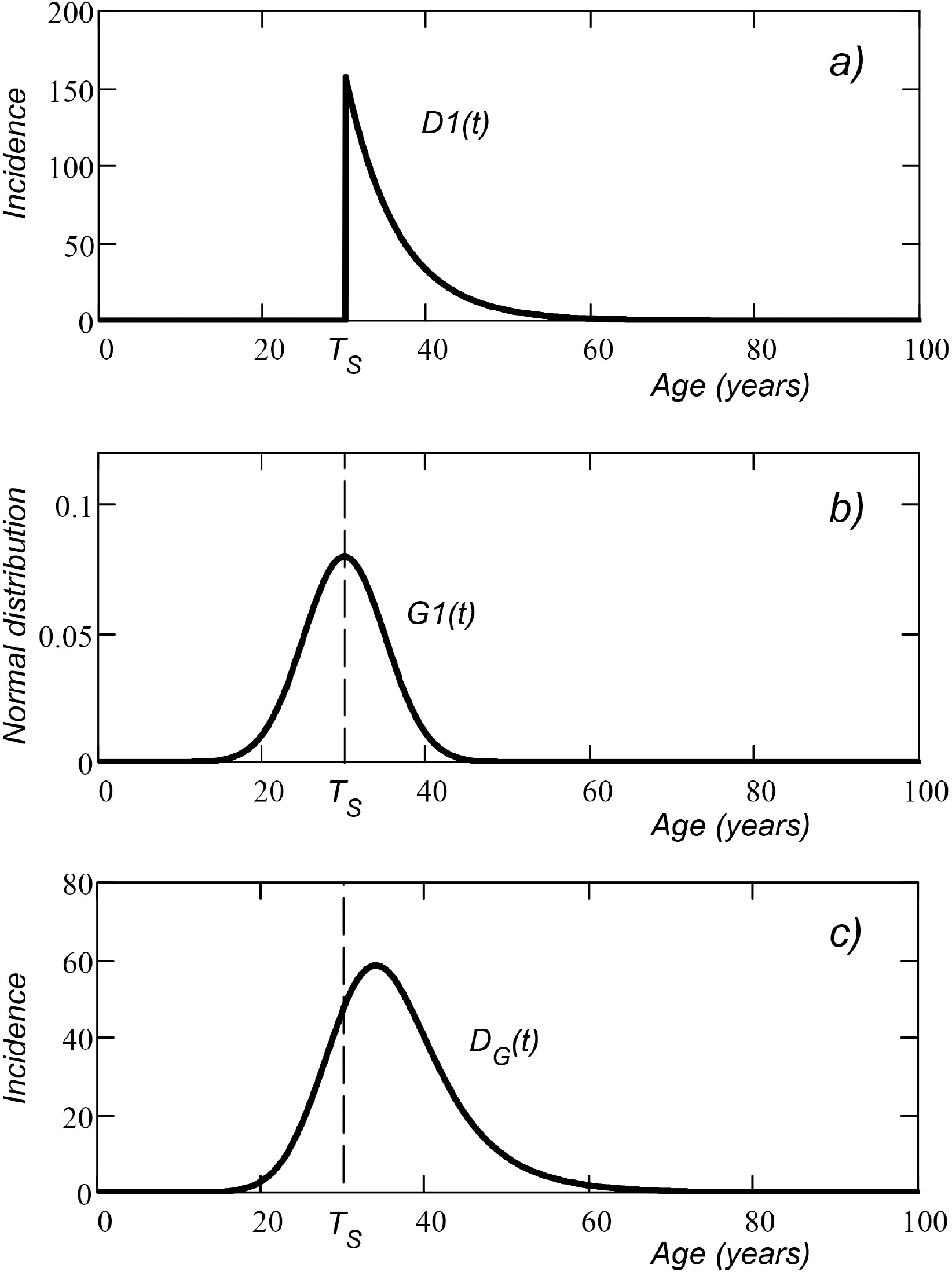
a) – the function of age distribution of cancers for a group of 1000 people with the same time *T s* = 30 (time of onset of carcinogenic effects); b) – Gaussian distribution of time *T s* in the group; c) – the function *D*_*G*_(*t*) of age distribution of cancers taking into account the normal distribution of time *T*_*S*_ in the group.

In this distribution, the age of *T*_*S*_ (the age at which the carcinogenic effect begins) is the same for all people. In real life, this is not the case. Some people have a slightly longer life expectancy than the average life expectancy in a group, while others have a slightly shorter life expectancy. The age of onset of carcinogenic effects is different for everyone.

Therefore, in reality, the sharp peak of the distribution is somewhat blurred along the time axis.

We assume that the age of onset of carcinogenic effects has a normal (Gaussian) distribution *G*1 (*t*) for the entire group (with the standard deviation *σ*):

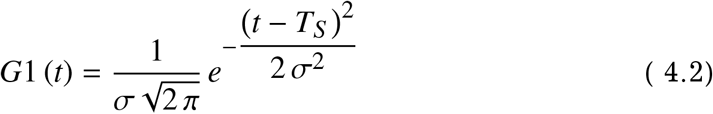

The normal distribution of the age of *T*_*S*_ in the group is shown in Fig. 4.1-b. The *σ* parameter (the Greek letter “sigma”) sets the width of the blurring of the graph. With a large *σ* the graph becomes wider, with a small value of *σ* it becomes narrower.

We can roughly estimate the value of the parameter *sigma* by the distribution of natural mortality in the population (mortality from old age). For example, if the average age of death in the population is 80 years, and the parameter *sigma* in the age distribution of mortality is 10, then for the age distribution of some event that occurs at the age of 40 (on average for a group) the value of *sigma* is approximately 5. Thus, the smaller the age of the event under consideration, the smaller the value of *sigma*. For zero age *sigma* = 0.

To find the age distribution *D*_*G*_(*t*) of cancers, taking into account a different time shift for each of the people in the population, we need to sum the distributions of *D*1(*t*) for all people, taking into account the age distribution of the time *T*_*S*_ in the group. Divide the time axis into segments of length *dt* and set the value of the age distribution on each segment as shown below.

For the first segment:

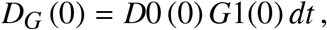

Here *D*0 (*t*) is the function *D*1(*t*) without time shift:

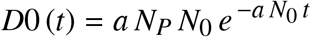

For the next moment in time *t* = *dt*:

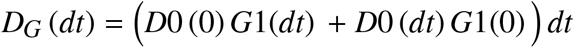

For the time *t* = 2*dt*:

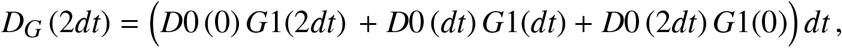

and so on:

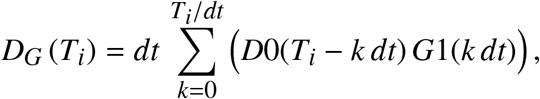

where *T*_*i*_ is *i dt*.

Now we can use the integral, counting, in the limit, *dt* → 0:

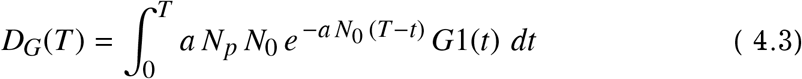

Here *T* and *t* is the same variable. We use different notation to avoid confusion with the limits of the integral when calculating.

This is the age distribution of cancers in a group of *N*_*P*_ people for a delayed carcinogenic event model. The age of onset of carcinogenic effects is *T*_*S*_. The distribution is built taking into account the normal distribution of the time *T*_*S*_ in the group (Fig. 4.1-b).

The graph of the function *D*_*G*_ (*t*) (age distribution of cancers in a group of 1000 people) is shown in Fig. 4.1-c.

This is the age distribution *D*1(*t*) shown in Fig. 4.1-a, which is “blurred” along the horizontal axis due to different times of the onset of carcinogenic effect for each person in the group. The average time of the onset of carcinogenic effect for the entire group is *T*_*S*_ (the time *T*_*S*_ is shown in the figure by the vertical dashed line).

We used the following parameters to plot the graph: *a* = 8 · 10^−8^; *N*_0_ = 2 · 10^6^; *N*_*P*_ = 1000; *T*_*S*_ = 30; *σ* = 5. Compare the graph in Fig. 4.1-a with the graph in Fig. 4.1-c (graphs have different scales of the vertical axis).

For this distribution, the normalization condition is satisfied:

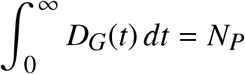

That is, the area under the curve of the function *D*_*G*_(*t*) is *N*_*P*_ (the size of group).

In this model of the age distribution of cancers, the *D*_*G*_(*t*) distribution graph always has the shape of a hill with a steep left slope and a gentler right slope. This is a characteristic feature of the model, due to which we cannot always use this model to approximate real age distributions.

## Part II Statistical analysis of age distributions of cancers

### 5. Practical datasets

The data presented in [6] were selected as datasets for statistical analysis. These are generalized data from the register of carcinogenic diseases in Belarus. Data are presented as the age distributions of the incidence of lung, stomach, colon and breast (mammary) cancer in women and the age distributions of lung, stomach, colon and prostate cancer in men.

Age distribution data are given per 100,000 people – this is convenient for numerical approximation. All datasets, except for data on the incidence of mammary cancer, are averaged for the period from 2006 to 2010. Mammary cancer is presented for 2010, without averaging statistics.

The datasets are presented in Fig. 6.1 (and also in Figures 7.1 and 8.1) as separate black square dots. Smooth curves on the graphs are approximating functions obtained separately for each model of age distribution (the parameters of the approximating functions are described in detail below in the corresponding sections).

**Figure 6.1.**
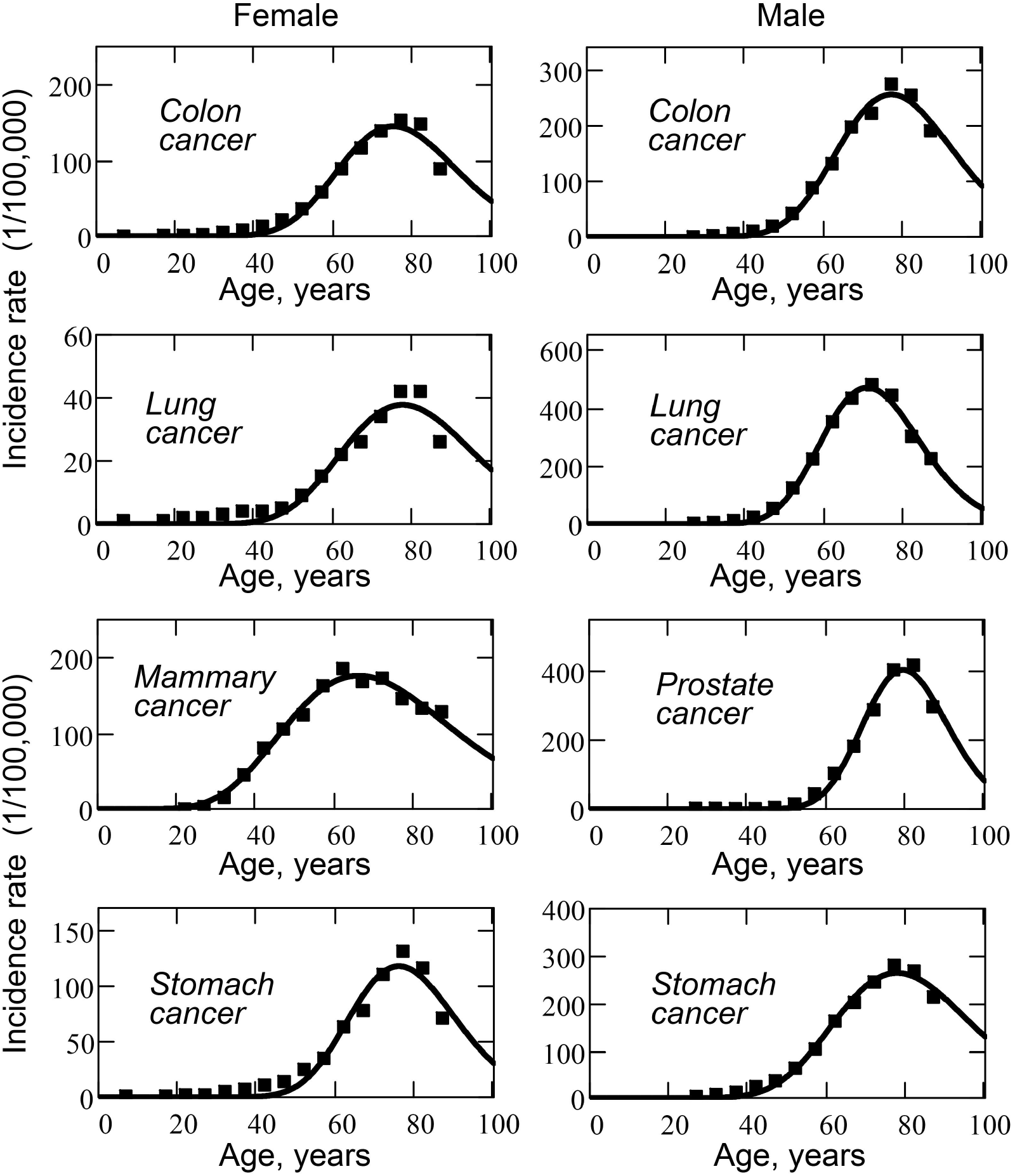
Age distributions of cancers per 100,000 people in women (left column) and men (right column), which are approximated by the function *Dz* (*t*) within a simple mutational model.

**Figure 7.1.**
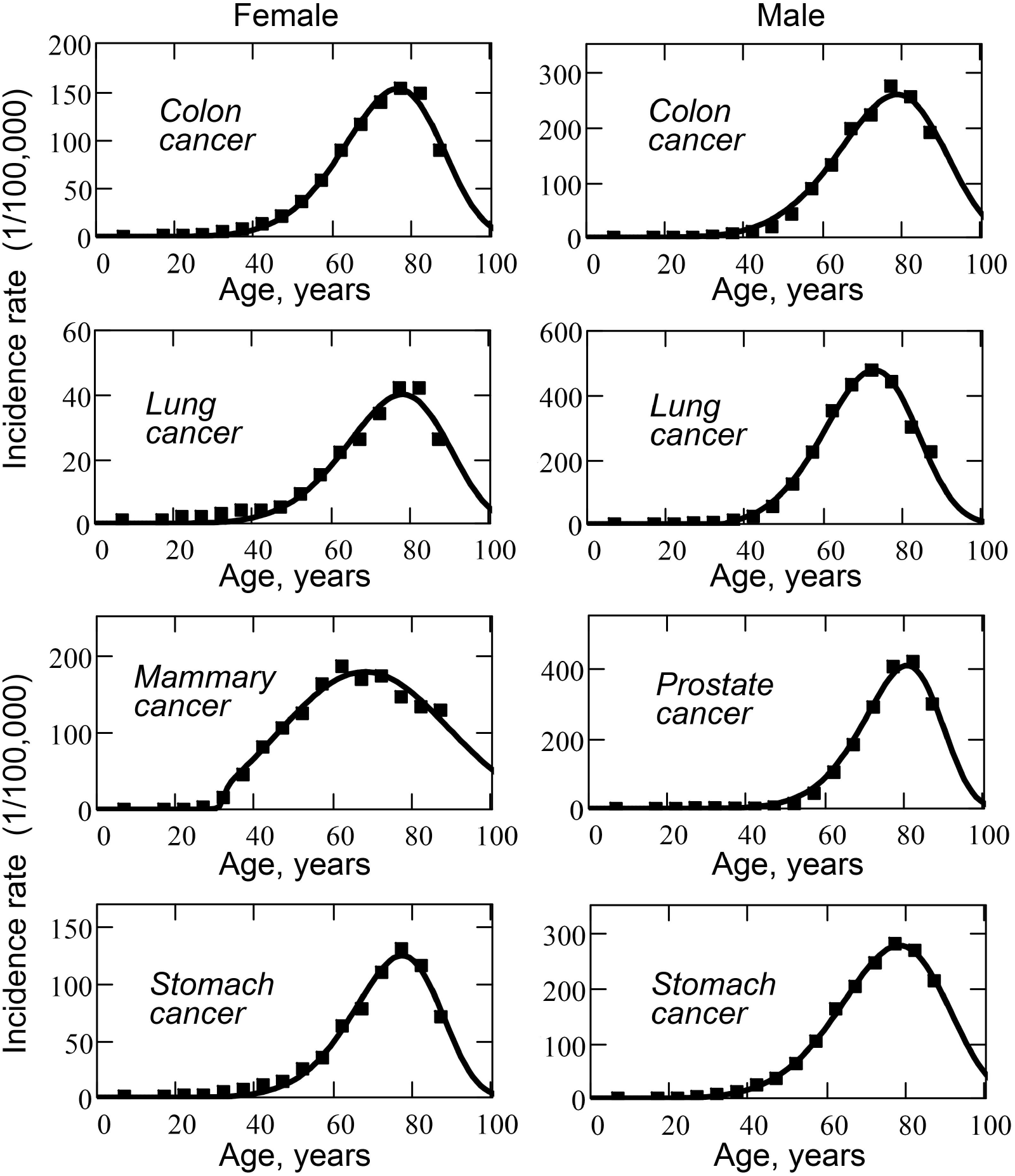
Age distributions of cancers in a group of 100,000 people in women (left column) and men (right column), which are approximated by the function *Dz* (*t*) within a complex mutational model.

**Figure 8.1.**
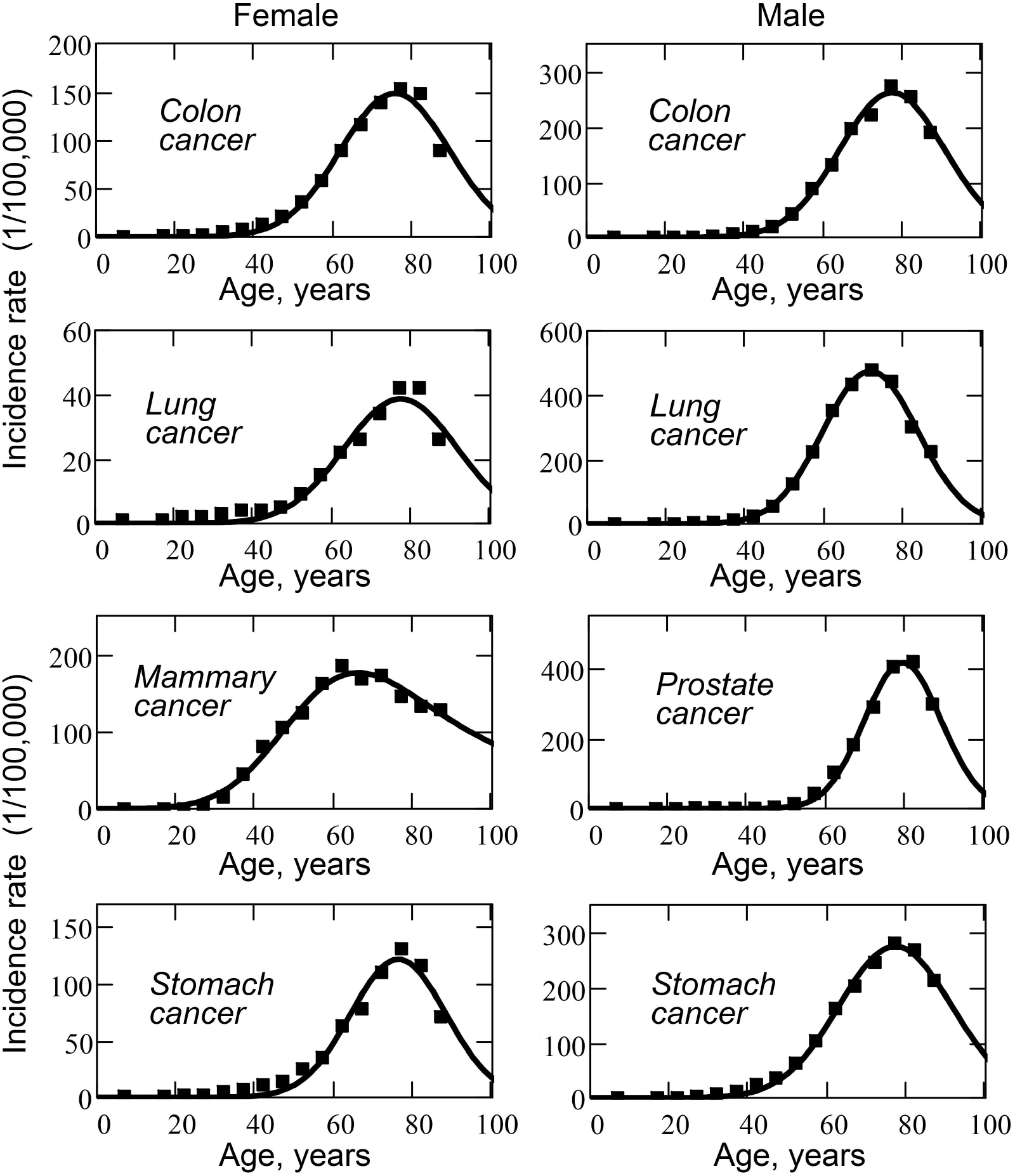
Age distributions of cancers in a group of 100,000 people in women (left column) and men (right column), which are approximated by the function *D*_*G*_ (*t*) within a model of delayed carcinogenic event (see Eq. 4.3).

The graphs in the left column of Fig. 6.1 show data for female cancers, the graphs in the right column show data for male cancers.

### 6. Statistical analysis based on a simple mutational model

A simple mutational model does not correspond much to reality, but (when optimizing the approximation parameters) it demonstrates the features of the approximating function behaviour, which are also characteristic of a more complex mutational model.

As an approximating function, we use the age distribution function *Dz* (*t*) (see Eq. 2.36):

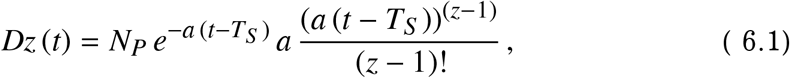

where *z* is the number of mutations in the cell, after which cancer necessarily begins.

Since cancer diagnosis is usually delayed in relation to the onset of the disease (cancer develops slowly, at the initial stage the cancer is practically invisible and is usually diagnosed after a few years), the formula includes the *T*_*S*_ lag time. The function is optimized for four parameters: *z, a, N*_*P*_, and *T*_*S*_. The least squares method is used to select a set of parameters for which the error function *Er* (*z, a, N*_*P*_, *T*_*S*_) has a minimum value:

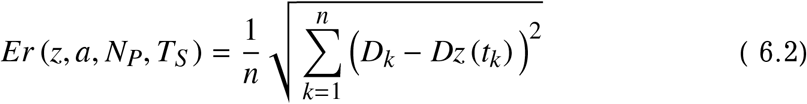

Here *t*_*k*_ is the age from the real dataset; *D*_*k*_ is the value of the age distribution from the real dataset for the age of *t*_*k*_; *Dz* (*t*_*k*_) is the theoretical value of the age distribution for the age *t*_*k*_.

The optimization of the function *Er* is carried out programmatically, by the Quasi-Newton method [7], for each fixed value of *z* (since the *z* parameter is a positive integer).

To compare the magnitude of the error for different age distributions, for the obtained (optimized) parameters, we calculate the value of the relative error *Er*_%_ (as a percentage of the maximum value *D*_*max*_ from the real dataset):

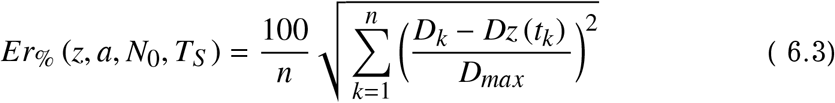

Here *D*_*max*_ is the maximum value of the age distribution function from the real dataset. The fixed maximum value is used due to the fact that in the real and theoretical distribution the points of the curves (points that are not close to the point of mathematical expectation) are very close to zero. This leads to a strong scatter of the relative error and the impossibility of comparing the optimization quality for different age distributions.

The obtained optimal parameters *z, a, N*_*P*_ and *T*_*S*_ and the values of the absolute and relative errors (the functions *Er* and *Er*_%_) are shown in Table 6.1 (separately for female and male cancers). In the bottom line of the table (for women and for men) the parameters, averaged over all types of cancer, are shown in bold.

**Table 6.1.**
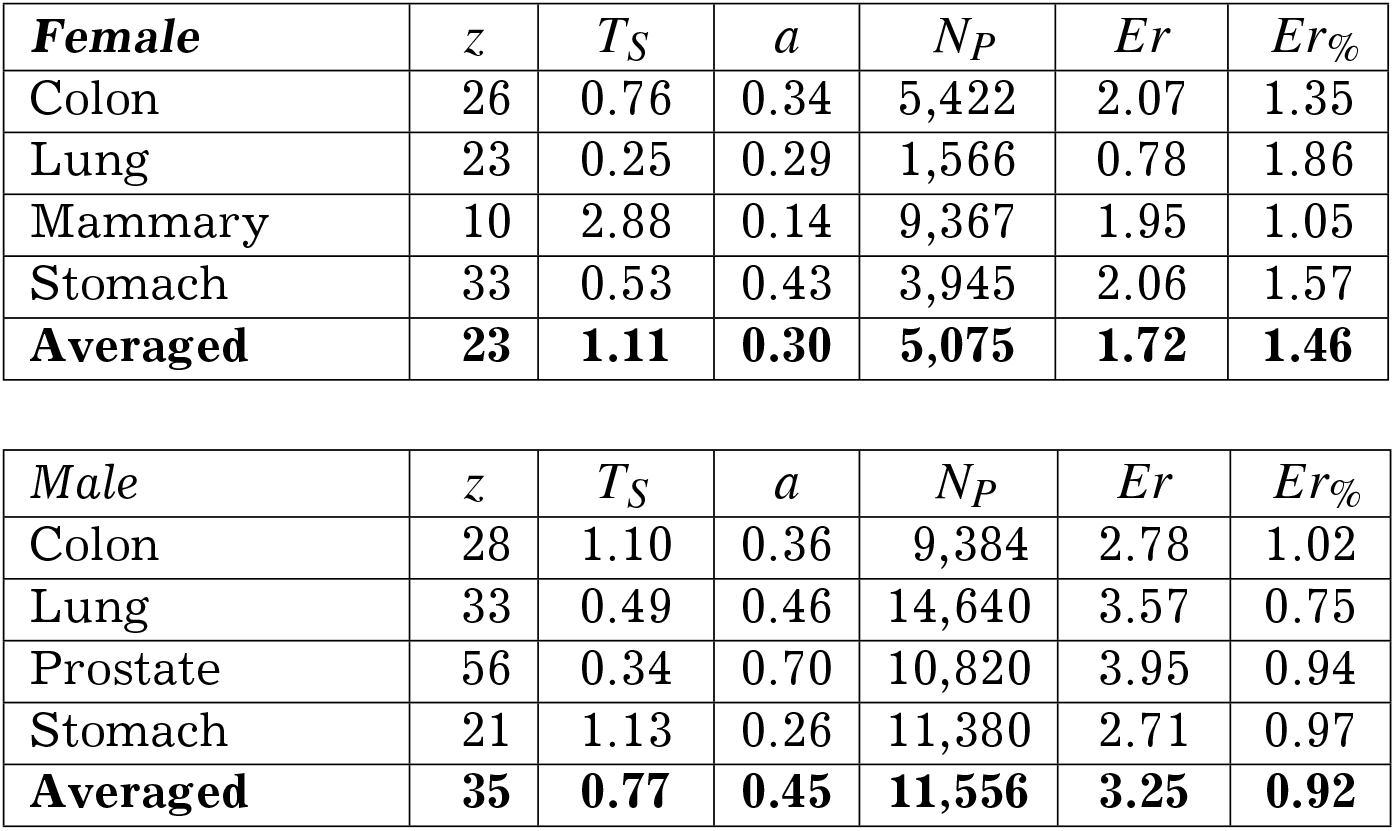
Fitting parameters for the simple mutation model.

The average number of mutations required to form a cancer is 23 mutations per cell for female cancers and 35 mutations per cell for male cancers. The average number of mutations per unit time is 0.30 mutations per year per cell in women and 0.45 mutations per year per cell in men.

The average (for all types of cancer) the number of people in the group forming this age distribution of cancer is 5,075 people for female cancer and 11,556 people for male cancer.

The approximating functions (the solid curved lines in graphs) corresponding to the optimal parameters of the function *Er* are shown in Fig. 6.1. The dark points in the graphs are datasets of the real age distribution of cancers.

### 7. Statistical analysis based on a complex mutational model

A complex mutational model takes into account the growth of the cell population and the transfer of mutations obtained by the parent cells to the daughter cells. Cell population growth occurs in two stages: at the initial stage the population grows exponentially, at the second stage the population has limited growth and the population size is gradually approaching a certain limit.

At the initial moment of time (*t* = 0) the population consists of one cell. Then the population increases. The final population size is given by the expression *R*_0_/*k*_0_. The *R*_0_ parameter is the value of the available energy (food) resource (which is available to the population per unit of time). The *R*_0_ parameter defines the final number of cells in the population at the time *t* = ∞.

We can estimate *N*_*G*_ = *R*_0_/*k*_0_ (the number of cells in a cell tissue of an adult organ) if we know the anatomical size of the human organ and the size of an individual cell [8].

In this work, we used the following estimates:

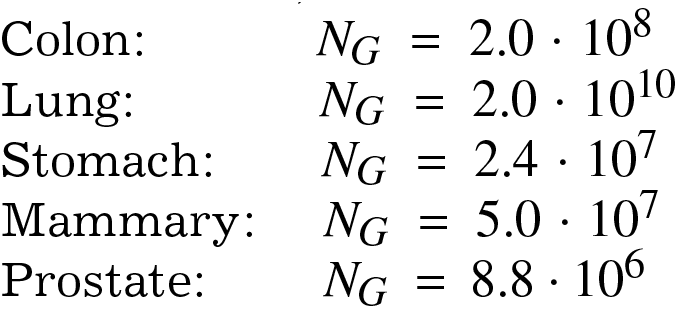

Parameter *R*_0_ is a constant environmental parameter. The parameters *k*_0_, *k*_1_, *k*_3_ are biological parameters of the cell population (see Section 2). These parameters are fixed and do not change during calculations.

The coefficients *k*_0_, *k*_1_, *k*_3_ are chosen so that the growth curve corresponds to our ideas about the growth rate of the human body. In our case, we believe that growth rate is maximum at the age of 15 years (*t*_1_ = 15). We use the following coefficient values: *k*_0_ = 0.5, *k*_1_ = 1. The *R*_0_ parameter is set so that the *R*_0_/*k*_0_ ratio would set the required number of cells in the cell population of the adult organ tissue. The *k*_3_ parameter is calculated from the condition *t*_1_ = 15.

For a complex mutational model, we use the function *Dz* (*t*) as an approximation function (fit function) for the age distribution of cancers (see Eq. 3.22):

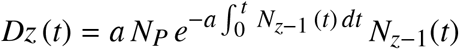

As before, in the simple mutation model, the delay parameter *T*_*S*_ is also optimized – we believe that cancer is diagnosed not at the time of the onset of the disease, but after a few years (after a time *T*_*S*_ since the onset of the disease).

That is, the *T*_*S*_ time must not be zero or negative.

We optimize the function by four parameters: *z, a, N*_*P*_, and *T*_*S*_. The least squares method is used to select a set of parameters for which the error function *Er* (*z, a, N*_*P*_, *T*_*S*_) has a minimum value:

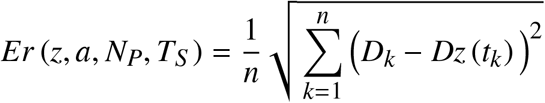

Here *t*_*k*_ is the age from the real dataset; *D*_*k*_ is the value of the age distribution from the real dataset for the age of *t*_*k*_; *Dz* (*t*_*k*_) is the theoretical value of the age distribution for the age *t*_*k*_.

To compare the magnitude of the error for different age distributions, for the obtained (optimized) parameters, we calculate the value of the relative error *Er*_%_ (as a percentage of the maximum value *D*_*max*_ from the real dataset):

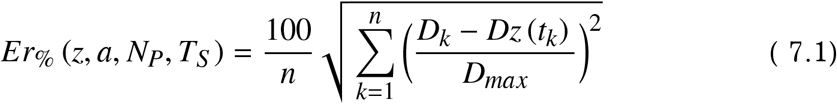

As before (for a simple mutation model), the optimization of the *Er* function is carried out programmatically, using the Quasi-Newton method for each fixed value of *z* (because the *z* parameter is a positive integer).

The obtained optimal parameters *z, a, N*_*P*_ and *T*_*S*_ and the values of the absolute and relative errors (the functions *Er* and *Er*_%_) are shown in Table 7.1 (separately for female and male cancers).

**Table 7.1.** Fitting parameters for the complex mutational model.

| **Female** | $z$ | $T_S$ | $a$ | $N_P$ | $Er$ | $Er\%$ |
| --- | --- | --- | --- | --- | --- | --- |
| Colon | 6 | 9.29 | $8.9 \cdot 10^{-1}$ | 4,741 | 1.09 | 0.71 |
| Lung | 6 | 8.56 | $7.3 \cdot 10^{-3}$ | 1,285 | 0.57 | 1.35 |
| Mammary | 3 | 17.00 | $7.6 \cdot 10^{-6}$ | 8,689 | 1.94 | 1.05 |
| Stomach | 7 | 6.54 | $2.2 \cdot 10^{-2}$ | 3,485 | 1.12 | 0.85 |
| **Averaged** | **5.50** | **10.35** | **$5.5 \cdot 10^{-1}$** | **4,550** | **1.18** | **0.99** |

| **Male** | $z$ | $T_S$ | $a$ | $N_P$ | $Er$ | $Er\%$ |
| --- | --- | --- | --- | --- | --- | --- |
| Colon | 6 | 12.86 | $1.1 \cdot 10^0$ | 8,063 | 1.46 | 0.53 |
| Lung | 5 | 20.44 | $3.3 \cdot 10^{-4}$ | 13,897 | 3.41 | 0.72 |
| Prostate | 8 | 15.49 | $2.1 \cdot 10^{-5}$ | 9,526 | 0.78 | 0.19 |
| Stomach | 6 | 5.21 | $4.5 \cdot 10^0$ | 9,348 | 0.93 | 0.33 |
| **Averaged** | **6.25** | **13.50** | **$5.3 \cdot 10^{-4}$** | **10,209** | **1.65** | **0.44** |

In the bottom line of the table (for women and for men) the parameters averaged over all types of cancer are shown in bold.

The average number of mutations required to form a cancer is 5.5 mutations per cell for female cancers and 6.25 mutations per cell for male cancers. The average number of mutations per unit time is 5.5 · 10 ^1^ mutations per year per cell for women and 5.3 · 10 ^4^ mutations per year per cell for men.

The approximating functions (the solid curved lines in graphs) corresponding to the optimal parameters of the function *Er* are shown in Fig. 7.1. The dark points in the graphs are datasets of the real age distribution of cancers.

### 8. Statistical analysis based on model of delayed carcinogenic event

In the model of delayed carcinogenic event (a single event with a time shift), to approximate the age distribution of cancers, we use the function *D*_*G*_ (*t*) (see Eq. 4.3):

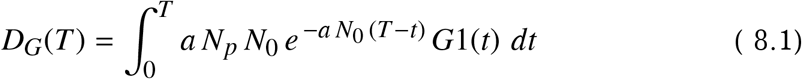

Here *G*1(*TS−t*) is a Gaussian distribution shifted to the right from zero by *TS* (see Eq. 4.2). Using the least squares method with a choice of parameters: *a, NP, TS, σ*, we minimize the error function (programmed coordinate descent method):

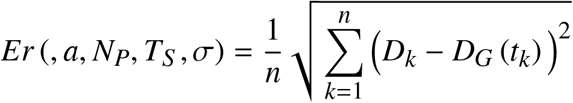

In this case, the *T*_*S*_ time is the sum of the average age of onset of carcinogenic effect and the time from the onset of cancer to diagnosis of the disease.

As before, we evaluate the function of relative error *Er*_%_ (as a percentage of the maximum value of *D*_*max*_ from the dataset of the real age distribution):

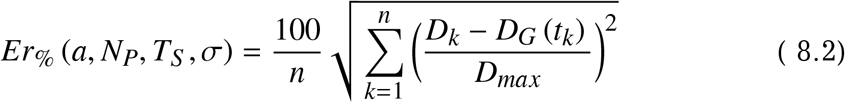

The obtained optimal parameters *a, N*_*P*_ and *T*_*S*_ and the absolute and relative error values (the functions *Er* and *Er*_%_) are shown in Table 8.1 separately for female and male cancers. In the bottom line of the table (for women and men), the parameters averaged over all types of cancer are shown in bold.

**Table 8.1.** Approximation parameters for the model of delayed carcinogenic event.

| ***Female*** | $N_P$ | $T_S$ | $a$ | $\sigma$ | $Er$ | $Er_{\%}$ |
| --- | --- | --- | --- | --- | --- | --- |
| Colon | 5,097 | 75.2 | $1.3 \cdot 10^{-8}$ | 13.7 | 1.62 | 1.06 |
| Lung | 1,381 | 75.4 | $2.6 \cdot 10^{-11}$ | 14.1 | 0.68 | 1.63 |
| Mammary | 11,378 | 50.7 | $5.8 \cdot 10^{-10}$ | 13.5 | 1.81 | 0.98 |
| Stomach | 3,661 | 75.9 | $1.3 \cdot 10^{-7}$ | 12.0 | 1.71 | 1.30 |
| **Averaged** | **5,379** | **69.3** | **$3.6 \cdot 10^{-8}$** | **13.3** | **1.46** | **1.24** |

| ***Male*** | $N_P$ | $T_S$ | $a$ | $\sigma$ | $Er$ | $Er_{\%}$ |
| --- | --- | --- | --- | --- | --- | --- |
| Colon | 8,718 | 74.6 | $1.8 \cdot 10^{-9}$ | 13.0 | 1.79 | 0.65 |
| Lung | 14,288 | 71.6 | $2.4 \cdot 10^{-9}$ | 12.0 | 2.26 | 0.47 |
| Prostate | 10,301 | 77.0 | $4.2 \cdot 10^{-8}$ | 9.6 | 2.48 | 0.59 |
| Stomach | 9,820 | 69.0 | $1.6 \cdot 10^{-8}$ | 14.0 | 1.70 | 0.61 |
| **Averaged** | **10,782** | **73.1** | **$1.6 \cdot 10^{-8}$** | **12.2** | **2.06** | **0.58** |

The average (for all types of cancer) number of people in the group forming a given age distribution of cancers is 5,379 people for female cancer and 10,782 people for male cancer. The average number of carcinogenic events (parameter *a*) per person per year is 3.6·10 ^−8^ events for women and 1.6·10 ^−8^ events for men. The average time *T*_*S*_ (the sum of the average age of onset of carcinogenic effect and the time from the onset of cancer to its diagnosis) is 69.3 years for women and 73.1 years for men.

The approximating functions (the solid curved lines in graphs) corresponding to the optimal parameters of the function *Er* are shown in Fig. 8.1. The dark points in the graphs are datasets of the real age distribution of cancers.

## Part III Discussion

### 9. Mutational models

Applying the mutational models considered in the work, we can determine (by applying the models to real age distributions) the following parameters of cancer:

– the number of mutations per cell required for cancer to appear (*z* parameter from Table 7.1);
– the average number of mutations per unit time (year) per one cell of the considered cell tissue (parameter *a* from Table 7.1);
– the time lag in medical diagnosis of cancer (the time interval between the occurrence of cancer in the cell and its detection), (*T*_*S*_ parameter from Table 7.1);
– the relative proportion of people susceptible (genetically predisposed) to this form of cancer (parameter *N*_*P*_/100, 000 from Table 7.1).

There are articles in which, based on real age distributions of cancers, the number of cell mutations required for cancer formation is estimated. For example, in [9], the Weibull distribution is used as a mathematical model, but the equations of the growth of the cell population are not taken into account.

In [10], real age distributions are approximated using 16 different probability distributions (including the Weibull distribution) – to determine which of these distributions gives the best approximation accuracy. The best of those considered is the Erlang distribution mentioned in Section 2.2.

But we do know that the Erlang distribution describes a simplified mutational model that is not accurate enough to match reality.

In our case, in order to understand which of the models gives the best results for approximating the real age distributions of cancers, we can compare the parameters of the relative mean square error (*Er*_%_ parameters in Tables 6.1 and 7.1) for the mutational models presented in this work.

For female and male cancers, a complex mutational model gives the best approximation results. Also, a complex mutational model is better than the delayed carcinogenic event model.

Another important parameter of the mutation model is the time lag between cancer formation and diagnosis.

However, when regression using a simple mutation model, this time delay is not detected (see *T*_*S*_ parameter in Table 6.1). Although this parameter is nonzero in the table, during optimization the program constantly reduces the time *T*_*S*_ to zero (and even to the area of negative values) in order to find the best approximation to a set of real data. Therefore, we write the last non-zero value of *T*_*S*_ into the table. Although, in theory, there should be a stable minimum of the error function to the left of the maximum of the practical data set. Medical practitioners usually estimate the diagnosis lag time (time from onset of cancer to diagnosis) at about 5-10 years.

In a complex mutational model, we find a delay in diagnosis in only half of the cases examined: colon cancer, lung cancer, prostate cancer in men, and breast cancer in women. (see table 7.1). The other forms of cancer considered do not have a time lag. As in the simple mutation model, we include in the table the last non-zero value of the *T*_*S*_ parameter, when the optimum of the error function was not found in the region of positive values of *T*_*S*_.

The table value *T*_*S*_ is not zero due to the fact that the number of mutations *z* is an integer, and, as an optimum, the table contains the maximum *z* for which *T*_*S*_ is not negative yet. If we allow negative values of *T*_*S*_, then for the error function we can get better values. This, perhaps, tells us that either the mutational hypothesis is wrong, or the presented mathematical model does not match reality well enough.

Quantitative analysis of mutational models shows results that are consistent with some of the concepts of the mutational cancer formation hypothesis. For example, for the female form of colon cancer, the size of the group of people providing the real age distribution of cancer is 4,741 people (for a complex mutational model). Although the real distribution is obtained for a group of 100,000 people. This may mean that only part of the group in question is predisposed to cancer (4,741 people of 100,000 – or 4.74%), and other people (95,259) don’t get cancer (probably for reasons of genetic resistance to the disease).

In other words, according to the data of the considered mutational model, cancer occurs only in a fraction of people who are probably genetically predisposed to the disease. (this is on average 4.55% of women and 10.21% of men for all types of cancer considered – see *N*_*P*_ parameter in Table 7.1). Cancer develops when one of the cells in the anatomical tissue receives a certain number of mutations. (on average, this is 5-6 mutations per cell – see *z* parameter in Table 7.1).

The disadvantage of the model is the large value of the parameter *a*, which is obtained by approximating the real age distribution for prostate cancer in men (*a* is the average number of mutations in a cell per unit time). The value of *a* parameter turns out to be incredibly high – 53,000 mutations per cell per year (or 145 mutations per cell per day). For example, the *a* parameter ranges from *a* = 7.6 · 10^−6^ for breast cancer in women to *a* = 2.1 · 10^5^ for prostate cancer in men. It turns out that in this model, the rate of formation of mutations in a cell strongly depends on the number of cells in the tissue of the organ under consideration – the fewer cells in the tissue, the greater the average number of mutations in a cell per unit of time.

This may also indirectly indicate that the model does not describe real age distributions well, and some refinement of the model is required.

### 10. Model of delayed carcinogenic event

The delayed carcinogenic event model was developed as an alternative to the mutational models. Using this model, we can determine (by applying the model to real age distributions) the following parameters of cancer:

– the average number of carcinogenic events per unit time (year) per one considered anatomical organ (*a* parameter from Table 8.1);
– the relative proportion of people susceptible (genetically predisposed) to this cancer (*N*_*P*_/100, 000 parameter from Table 8.1).
– the sum of the time of the onset of carcinogenic effects and the time of delay in cancer diagnosis (*T*_*S*_ parameter from Table 8.1);
– the degree of non-simultaneity of the onset of carcinogenic effects in different people in the group (*σ* parameter from Table 8.1);

The model of a delayed carcinogenic event (with the same number of parameters of the approximating function) gives, on average, a smaller value of the relative root-mean-square error (parameter *Er*_%_) in comparison with the simple mutational model, but loses in terms of the magnitude of the error in comparison with the complex mutational model (compare the average values of *Er*_%_ in tables 6.1, 7.1 and 8.1).

As well as for mutational models, we can quantify the main parameters of the mathematical model by comparing the parameters with the real age distributions of cancers.

For example, for prostate cancer, the size of the group of people that provides the real age distribution of cancers is 10,301 people. This means that only a given number of men will get prostate cancer in their lives. This can be explained by a hereditary predisposition to the disease. That is, 10,3 % of men are prone to prostate cancer.

The disadvantage of the model is the impossibility to separate (from the sum of two values) the delay time of cancer diagnosis (the time between the onset of cancer and the diagnosis of the disease). The time *T*_*S*_ in the model is the sum of the time of the onset of carcinogenic effect and time between the onset of cancer and the diagnosis of the disease.

The model does not tell us anything about the cause of cancer, but, nevertheless, it allows us to estimate some parameters of the carcinogenic damaging effect. In particular, we can estimate the average number of carcinogenic events per cell per unit of time. This is the *a* parameter from Table 8.1.

In any case, in order to further explore the models presented in this article, we need to match the models with a lot of real data on the age distribution of cancers.

Saint-Petersburg

2021

## References

[1] Loeb L.A., Loeb K.R., Anderson J.P. Multiple mutations and cancer PNAS, Vol. 100, No. 3, pp. 776–781, February 2003. https://doi.org/10.1073/pnas.0334858100

[2] Knudson A.G., JR. Mutation and Cancer: Statistical Study of Retinoblastoma. Proc. Nat. Acad. Sci.USA, Vol. 68, No., pp. 820–823, April 1971. https://doi.org/10.1073/pnas.68.4.820

[3] Tetearing A.N. Numerical Model of Fishery Impact on Population Levels of Landlocked Salmon in Lake Onego (Russia) 2020, https://dx.doi.org/10.2139/ssrn.3443512

[4] Erlang A.K. Telephone waiting times. Matematisk Tidsskrift. B, Vol. 31, p.25, Copenhagen, 1920

[5] Tetearing A.N. Theory of populations. SSO Foundation, Moscow,2012.

[6] Antipova S.I., Antipov V.V., Shebeko N.G. Gender problems of oncology in Belarus. Medical News Journal, No. 3., Minsk, 2013 (in Russian).

[7] Gill P.E., Murray W., Wright M.H. Practical optimization. Academic Press, London, 1981.

[8] Great Medical Encyclopedia. Edition 3, Moscow, 1974-1989 (in Russian).

[9] Calabrese P., Tavare S., Shibata D. Pretumor Progression. Clonal Evolution of Human Stem Cell Populations. American Journal of Pathology, Vol.164, Issue 4, pp. 1337–1346, 2004, https://doi.org/10.1038/s41598-017-12448-7

[10] Belikov A.V. The number of key carcinogenic events can be predicted from cancer incidence. Scientific Reports, 7, (12170), 2017, https://doi.org/10.1038/s41598-017-12448-7

